# Topological screen identifies hundreds of Cp190 and CTCF dependent *Drosophila* chromatin insulator elements

**DOI:** 10.1101/2022.07.18.500462

**Authors:** Tatyana G. Kahn, Mikhail Savitsky, Chikuan Kuong, Caroline Jacquer, Giacomo Cavalli, Jia-Ming Chang, Yuri B. Schwartz

## Abstract

*Drosophila* insulators were the first DNA elements discovered to regulate gene expression by delimiting chromatin contacts. Remarkably, it is still unclear how many of them exist in the *Drosophila* genome and whether they have a pervasive impact on the genome folding. Contrary to vertebrates, there is no evidence that fly insulators block cohesin-mediated chromatin loop extrusion. Therefore, their mechanism of action remains an open question. To bridge these gaps, we mapped genomic contacts, transcriptomes and binding landscapes of insulator associated proteins in *Drosophila* cells deficient for CTCF and Cp190. With this approach, we discovered hundreds of chromatin insulator elements. Their study indicates that *Drosophila* insulators play a minor role in shaping the overall chromosome folding patterns but impact chromatin contacts locally at many individual loci. Our observations argue that Cp190 promotes co-binding of other insulator proteins and that the model, where *Drosophila* insulators block chromatin contacts by forming loops, needs revision. The extended catalogue of insulator elements presented here provides a significant new resource to study mechanisms that shape the folding of eukaryotic genomes.

## Introduction

Eukaryotic chromosomes are extensively folded in order to fit inside micrometre-size cell nuclei. The degree of folding varies between different regions of the chromosome and the specific folding patterns vary from cell to cell. Nevertheless, high-resolution imaging (*1–6*) and high-throughput Chromosome Conformation Capture (Hi-C) assays (*7–9*) indicate that certain spatial conformations appear more frequently or persist longer than others. Averaged over large populations of cells, these conformations appear as sub-megabase long chromatin regions, often referred to as topologically associating domains (TADs). Any two loci situated within such domain are more frequently in proximity compared to any two loci positioned in the two neighbouring TADs (*10, 11*).

What mechanisms cause the folding biases? To what extent do these biases influence gene expression? Both questions remain subject of debate. Electrostatic interaction between nucleosomes, DNA supercoiling, chromatin loop extrusion by the cohesin complexes and chromatin insulator elements were all proposed to play a role in shaping genome folding (*12, 13*). Here, we will focus on chromatin insulators as they seem to have evolved for regulation of gene expression.

These elements were first discovered in *Drosophila melanogaster* (*14–16*) but later identified in several developmental genes of flies and vertebrates (*17–27*). Based on transgenic experiments in *Drosophila*, it was proposed that chromatin insulators bias chromatin folding by interacting with each other (*28, 29*). In this view, chromatin loops, formed by two or more interacting insulator elements, compete with contacts between chromatin sites inside and outside the loop. It was hypothesized that insulator-binding proteins equipped with protein-protein interaction domains hold the insulator elements together. The “bridging” proteins are recruited to insulator elements by auxiliary sequence specific DNA binding proteins.

Consistently, a number of sequence specific DNA binding proteins (CTCF, Su(Hw), Ibf1, Ibf2, Pita, ZIPIC, BEAF-32) and two candidate “bridging” proteins (Cp190 or Mod(mdg4)) were implicated in *Drosophila* insulator function by genetic and biochemical screens (reviewed in: (*30*)). From those only one, CTCF, has a clear orthologue in vertebrates. As would be expected from the model, mammalian CTCF is frequently found at bases of chromatin loops detected by Hi-C. However, in this case, the correspondence is attributed to CTCF acting as a barrier for cohesin complexes extruding chromatin loops (*31, 32*). This process does not involve direct interactions between the two CTCF bound elements. Contrary to vertebrates, there is no evidence that *Drosophila* insulators block cohesin-mediated chromatin loop extrusion. Furthermore, Hi-C experiments in fly embryos and cultured cells identified very few chromatin loops that may be linked to insulator protein binding sites (*33–35*). Thus, whether the “looping model” applies to *Drosophila* insulators remains an open question.

How many fly genes are equipped with insulator elements is another question proved difficult to address. Although *Drosophila* was the first multicellular organism where genomic distributions of multiple insulator proteins became available (*36–38*) this did not solve the problem. It turned out that the binding of individual proteins and even their combinations is a poor predictor of whether a site contains a functional insulator element (*37, 39*). Therefore, only a couple of dozen *Drosophila* insulator elements have been characterized to date by transgenic assays that tested the blocking of enhancer-promoter communications or the spreading of a histone modification from a site tethering a histone methyltransferase (reviewed in: (*30, 40*)).

To close this gap, we undertook the parallel mapping of genomic contacts, transcriptomes and genomic binding landscapes of insulator binding proteins in cultured cells custom derived from *Drosophila* embryos homozygous for loss-of-function mutation in *CTCF* or *Cp190* genes. With this approach, we discovered hundreds of chromatin insulator elements. Their study indicates that chromatin insulators impact chromatin contacts locally at many individual loci and argues against the model where *Drosophila* insulators block chromatin contacts by forming insulator-insulator contacts.

## Results

To derive CTCF and Cp190 deficient cells, we used the Ras^V12^ transformation approach (*41*) and embryos homozygous for *CTCF^y+1^* and *Cp190^3^* mutations (Figure S1). The *CTCF^y+1^* allele is a 3.3kb deletion that removes the entire open reading frame of the *CTCF* gene and produces no protein (*39, 42*). The *Cp190^3^*is a point mutation that results in premature translation termination at position Q61 and a short non-functional product (*43*). Although CTCF and Cp190 proteins are essential for fly viability, the mutant cells are viable and proliferate in culture. We derived two *Cp190^3^*and two *CTCF^y+1^* mutant cell lines. In all cases, the presence of the corresponding mutation was confirmed by PCR-genotyping (Figure S1B) and sequencing (Figure S1D) so we focused our analyses on Cp190-deficient line CP-R6 and CTCF-deficient line CTCF 19.7-1c, hereafter referred to as Cp190 knock-out (Cp190-KO) and CTCF knock-out (CTCF-KO) cells. Western blot analyses detected no Cp190 in the Cp190-KO or CTCF in the CTCF-KO cell lines (Figure 1A) and the ChIP-qPCR analysis of previously characterized binding sites (*39, 44, 45*) confirmed the loss of Cp190 and CTCF from the chromatin of the corresponding mutant cells (Figure 1B). Western-blot assay indicates that the loss of CTCF does not cause significant reduction in the overall Cp190 level and, inversely, the CTCF level is not affected by the *Cp190^3^* mutation (Figure 1A). Similarly, the ablation of Cp190 or CTCF does not alter bulk levels of several other key insulator proteins tested (Figure S2). From this, we concluded that our mutant cell lines are a valuable system that provides large quantities of material to interrogate specific roles of Cp190 and CTCF in the three-dimensional genome organization.

**Figure 1.**
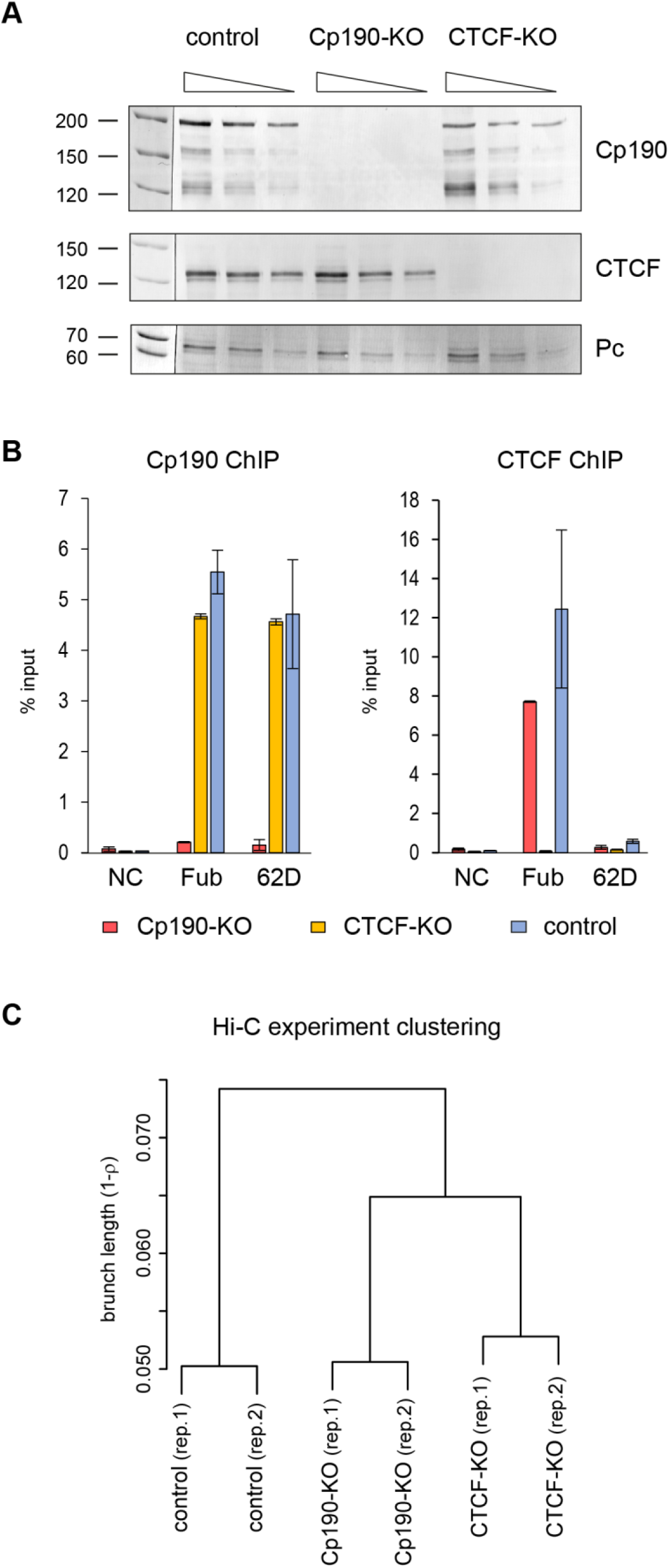
Characterization of Cp190- and CTCF-deficient cultured cell lines. **A.** Two-fold dilutions of total nuclear protein from wild type (Ras 3), Cp190-KO (CP-R6) and CTCF-KO (CTCF 19.7-1c) cells were analysed by western-blot with antibodies against Cp190, CTCF and Pc (loading control). Additional loading control, coomassie stained gel of corresponding total nuclear protein samples, is shown on Figure S2. Positions of molecular weight markers (in kDa) are indicated to the left. **B**. Chromatin Immunoprecipitation coupled to quantitative PCR (ChIP-qPCR) demonstrates that Cp190, normally present at *Fub* and *62D* insulators, is no longer detectable at these elements in Cp190-KO cells. Similarly, immunoprecipitation of *Fub* by anti-CTCF antibodies is abolished by CTCF knock-out. Histograms show the average of two independent ChIP-qPCR experiments with whiskers indicating the scatter between individual measurements. An intergenic region from chromosome 3L that does not bind any insulator proteins was used as negative control (NC). **C.** Hierarchical clustering of Hi-C experiments based on pairwise Spearman’s correlation coefficients (average ρ for the group). To suppress spurious experimental noise, contacts within individual 40kb bins (the diagonal of contact matrix) as well as contacts between bins separated by more than 1.6Mb were excluded from calculations. For bins equal or larger than 40kb, the clustering is robust to parameter changes (Figure S4).

### Cp190 loss disrupts topological organization of the homeotic gene cluster

How does the loss of Cp190 or CTCF affect the three-dimensional conformation of the genome? To address this question, we used the Hi-C assay (*9*) to map chromatin contacts in CTCF-KO, Cp190-KO and Ras^V12^ transformed but otherwise wild-type cells (Ras3, control) (*46*). After removing sequencing reads corresponding to circularized, unligated or non-digested fragments we detected from 44,406,664 to 81,238,455 chromatin contacts per each replicate and each genetic condition (for detailed statistics see Tables S1, S2). Assigned to genomic segments of fixed size (Hi-C bins) contact frequencies measured in replicate experiments are highly correlated. Correlation progressively increases when bins of larger size are analysed (from ρ = 0.81-0.88 for small 5kb bin to ρ = 0.98-0.99 for 160kb bin) and, overall, indicates that our Hi-C assay is reproducible. The contact frequencies remain strongly correlated when compared between different cell lines (Figure S3A) and, visualized at chromosome arm scale, contact maps of all three cell lines appear similar (Figure S3B). This argues that Cp190 or CTCF ablation does not grossly disrupt genome folding. Nevertheless, hierarchical clustering indicates that contact patterns in Cp190-KO and CTCF-KO cells are measurably distinct from those of the control cells (Figures 1C, S4).

To understand these differences, we started with a close inspection of the bithorax cluster of homeotic genes. The bithorax complex consists of three genes *Ubx*, *abd-A* and *Abd-B*, (Figure 2A) which encode transcription factors responsible for segmental identity of the abdomen and posterior thorax (*47*). Correct segment-specific expression of *Ubx*, *abd-A* and *Abd-B* is achieved by coordinated action of distal enhancers and Polycomb Response Elements (PREs). These enhancers and PREs are clustered in genetically defined domains (Figure 2A). The domains *abx/bx* and *bxd/pbx* control expression of *Ubx*. Series of *infra-abdominal* (*iab*) domains control expression of *abd-A* (*iab-2*, *iab-3*, *iab-4*) and *Abd-B* (*iab-5*, *iab-6*, *iab-7*, *iab-8*) (*48–50*). Five known insulator elements (*Fub*, *Mcp*, *Fab-6*, *Fab-7*, *Fab-8*) are required to ensure that the enhancers and PREs activate or repress the correct genes in correct body segments (*17, 19, 25, 51*). Of these, *Fub* is exceptionally robust in that it can block enhancer-promoter interactions regardless of its location in the genome (*39*). The *Fub* insulator is required to prevent erroneous activation of *abd-A* by the *Ubx* enhancers (*19, 52*) and likely has emerged after evolutionarily recent relocation of the *Ubx* gene from the *Antennapedia* gene cluster to the bithorax complex (*19*).

**Figure 2.**
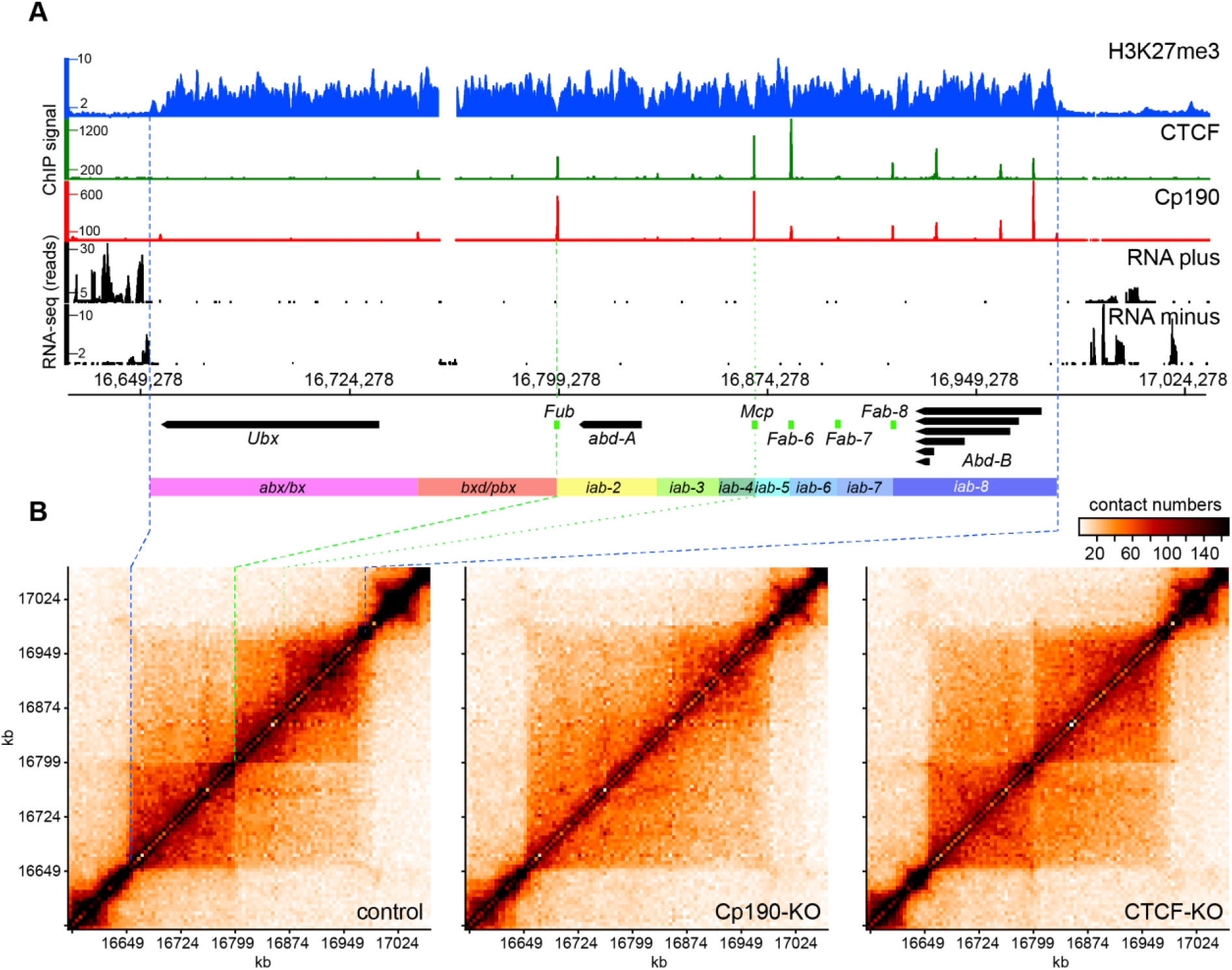
Organization and chromatin topology of the bithorax complex. **A.** Genomic organization of the bithorax complex. ChIP-on-chip profiles of H3K27me3 in ML-DmBG3-c2 cells from (*53*) displayed as IP/Input ratio, ChIP-seq profiles for Cp190 and CTCF in control cells (this study, displayed as number of sequencing reads per position per million of total reads) and RNA-seq profiles from control cells (displayed separately for each DNA strand as number of sequencing reads per position) are shown above the coordinate scale (*dm6*, 2014 genome release). The positions of main alternative transcripts for *Ubx*, *abd-A* and *Abd-B* genes are shown as thick arrows pointing in direction of transcription. Note, that transcripts flanking the bithorax complex genes are omitted for clarity. The positions of genetically defined insulator elements are indicated with green boxes. Regulatory domains are indicated as coloured rectangles. **B.** Chromatin contacts within the bithorax complex of the control, Cp190-KO and CTCF-KO cells. The contacts measured by individual Hi-C experiments were assigned to 5kb bins and normalized by iterative correction (*89*). The data from replicate experiments were combined and plotted with *gcMapExplorer* software (*80*). The correspondence between the edges of the H3K27me3 domain in **A** and **B** is shown with blue dashed lines. The green dashed line indicates the position of the *Fub* insulator element and the green dotted line shows the location of the *Mcp* insulator element.

In the control cells, *Ubx*, *abd-A* and *Abd-B* are repressed by Polycomb mechanisms (*46*). Sequencing of total RNA (RNA-seq) confirms that in these cells all three genes are transcriptionally inactive (Figure 2A). As illustrated by Figure 2B, in the control Ras3 cells, the bithorax complex is contained within a TAD whose borders match the borders of the chromatin domain enriched in histone H3 tri-methylated at Lysine 27 (H3K27me3) (*53, 54*). This domain is further split into two obvious large sub-domains at position precisely matching that of the *Fub* insulator element (Figure 2A, B). The outstanding enhancer blocking activity of *Fub* requires Cp190 but not CTCF (*39*). This is because, in addition to CTCF, *Fub* contains recognition sequences for other DNA binding proteins including Su(Hw). The latter directly interacts with Cp190 and can tether Cp190 to *Fub* even when CTCF is absent (*52*). In perfect agreement with genetic and molecular data, in Cp190-KO cells, but not in the CTCF-KO cells, the topological boundary between the *Ubx* and *abd-A* genes disappears (Figure 2B).

In addition, two distinct density clouds of chromatin contacts within *abd-A* and *Abd-B* genes and their regulatory regions are clearly visible in the contact map of the control cells (Figure 2B, the upper right corner of the bithorax complex TAD). The clouds segregate at approximately the site of the *Mcp* insulator element (even though our Hi-C assay does not single out *Mcp* as a clear-cut boundary) and they are no longer visible in the contact maps from the Cp190-KO and CTCF-KO cells.

Three conclusions follow from the observations above. First, our experimental system is sufficiently sensitive and accurate to detect topological changes around robust enhancer blocking insulators. However, it may miss those associated with weaker elements. Second, the *Fub* insulator is capable to limit chromatin contacts when the entire bithorax complex is transcriptionally inactive and repressed by Polycomb mechanisms. Finally, our observations suggest that a systematic screen for Cp190 and CTCF binding sites that limit chromatin contacts in the control cells but lose this ability in Cp190-KO and/or CTCF-KO cells may serve as genome-wide approach to discover robust insulator elements.

### Genome-wide survey of *Drosophila* insulator elements

Insulator proteins bind the genome in distinct combinations (*37, 38*) and some of these co-binding combinations correlate with enhancer-blocking ability. Nevertheless, it is not possible to predict *Drosophila* insulator elements from genome-wide mapping data alone (*37*). As a first step to bridge this gap, we tested whether our strategy detects known insulator elements. About thirty *Drosophila* insulator elements, including those from the bithorax complex, have been identified by genetic assays to date. Only one of them, the *gypsy*-like *62D* insulator element from the intergenic region between the *ACXD* and *CG32301* genes, has a robust position independent enhancer blocking capacity comparable to that of the *Fub* (*44, 55*). *62D* insulator binds Su(Hw), Cp190 and Mod(mdg4) but not CTCF (Figure 1B, (*44, 55*)). Consistently, in the control and CTCF-KO cells, the position of the *62D* insulator element coincides with a point of reduced contact crossing, which is alleviated in Cp190-KO cells (Figure S5).

Encouraged by this observation, we mapped genomic binding of Cp190, CTCF and several other key insulator proteins: Su(Hw), Mod(mdg4) and Ibf1 in the control, CTCF-KO and Cp190-KO cells using ChIP coupled to sequencing of the precipitated DNA (ChIP-seq). We performed two ChIP-seq experiments for each genetic background using independently prepared chromatins and had the DNA from corresponding chromatin input materials sequenced to control for possible sample processing biases. We then used Model-based Analysis of ChIP-Seq (MACS) algorithm (*56*) to identify genomic regions significantly enriched by immunoprecipitation with individual antibodies in the control cells and compared their genomic positions pairwise as illustrated in Figure 3. This approach grouped all enriched regions according to 26 common co-binding patterns (co-binding classes), which we designated with combinations of single letters that represent individual insulator proteins. For example, regions co-bound by **<u>M</u>**od(mdg4), **<u>C</u>**p190, **<u>I</u>**bf1, CTC**<u>F</u>** and **<u>S</u>**u(Hw) were designated as MCIFS.

**Figure 3.**
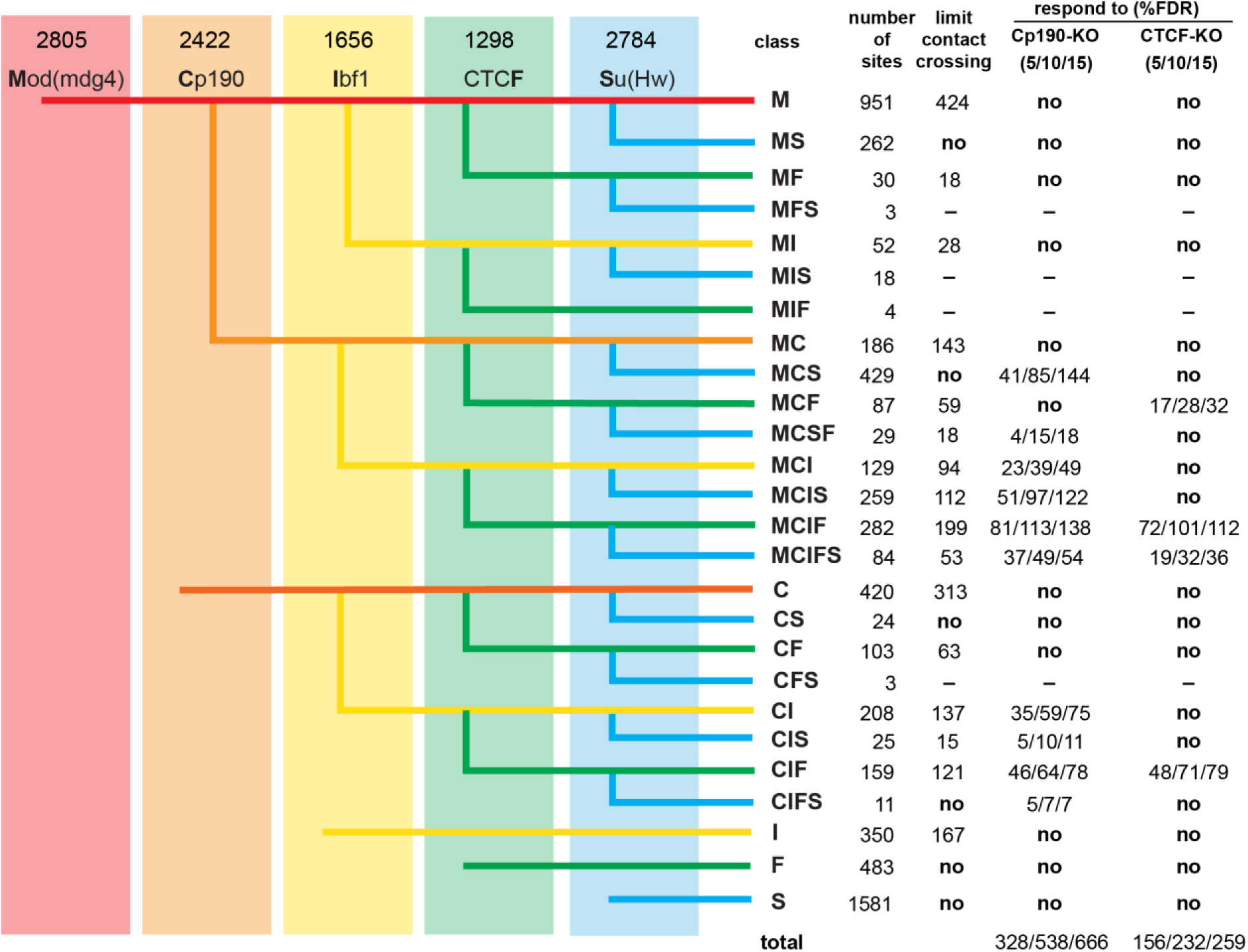
Classes of insulator protein binding regions. Genomic positions of regions enriched by immunoprecipitation of the chromatin from control cells (numbers indicated above each antibody) were compared pairwise in order from left to right. Thus, regions enriched by ChIP with antibodies against Mod(mdg4) were checked for overlap with regions enriched with antibodies against Cp190. The resulting three groups, i.e. bound by both Mod(mdg4) and Cp190, bound by just Mod(mdg4) or bound by just Cp190, were further compared to regions enriched by ChIP with antibodies against Ibf and so on. The resulted co-binding classes were designated with combinations of single letters representing individual insulator proteins present. Columns to the right indicate the number of regions within each class. For classes whose fraction of regions above the corresponding threshold significantly exceeds that of the control, the number of regions that limit contact crossing (γ > 75% of that for the control regions) or the number of regions that display increase in the contact crossing (Δγ ≤ FDR) upon Cp190-KO or CTCF-KO are shown further to the right. MFS, MIS, MIF and CFS sites were too few and, therefore, excluded from analyses.

How many regions in each co-binding class restrict chromosomal contacts? What fraction of those cease to limit contacts in Cp190-KO and/or CTCF-KO cells? To address these questions, we used Hi-C measurements to calculate the propensity of the chromatin contacts to cross insulator protein bound regions in the three cell lines. The frequency with which interaction between any two chromosomal sites is captured by Hi-C decays exponentially with increasing genomic distance between the sites (*9, 57, 58*). The global decay in contact frequency as a function of genomic distance is similar for all chromosome arms and can be approximated by a power law with scaling exponent derived from Hi-C measurements (*9, 32*). For most chromosomal sites, the frequency of pairwise interactions follows the global decay model and is, therefore, predictable from genomic distance between them. However, if two sites are separated by a region that limits chromosomal contacts, the observed interaction frequency is lower than that predicted by the global decay model. The prediction is improved by fitting a *distance-scaling factor* (γ) to each restriction fragment assayed in Hi-C (*9*). In this approach, regions that limit chromosomal contacts are assigned high distance-scaling factors. Using a computational pipeline developed by Yaffe and colleagues (*9*), we calculated γ for restriction fragments participating in the Hi-C assay and assigned each insulator protein bound region the highest γ from all restriction fragments overlapped by that region.

As illustrated by Figure 4A, some classes of insulator protein bound regions tend to limit contact crossing (tend to have high γ), while others are no different from randomly chosen control regions that do not bind insulator proteins. As noted previously (*37, 38*), the classes that tend to limit contact crossing tend co-bind multiple insulator proteins. However, the simple binding of large protein complexes does not explain the effect. For example, Polycomb Response Elements (PREs), which bind mega Dalton-size Polycomb complexes, have a distribution of γ similar to that of the control regions (Figure 4A). Consistent with previous transgenic enhancer-blocking tests (*37*), regions that bind CTCF but no Mod(mdg4) or Cp190 (“standalone” F class) do not limit contact crossing and have low γ. This indicates that *Drosophila* CTCF requires additional partners to affect the chromatin topology.

**Figure 4.**
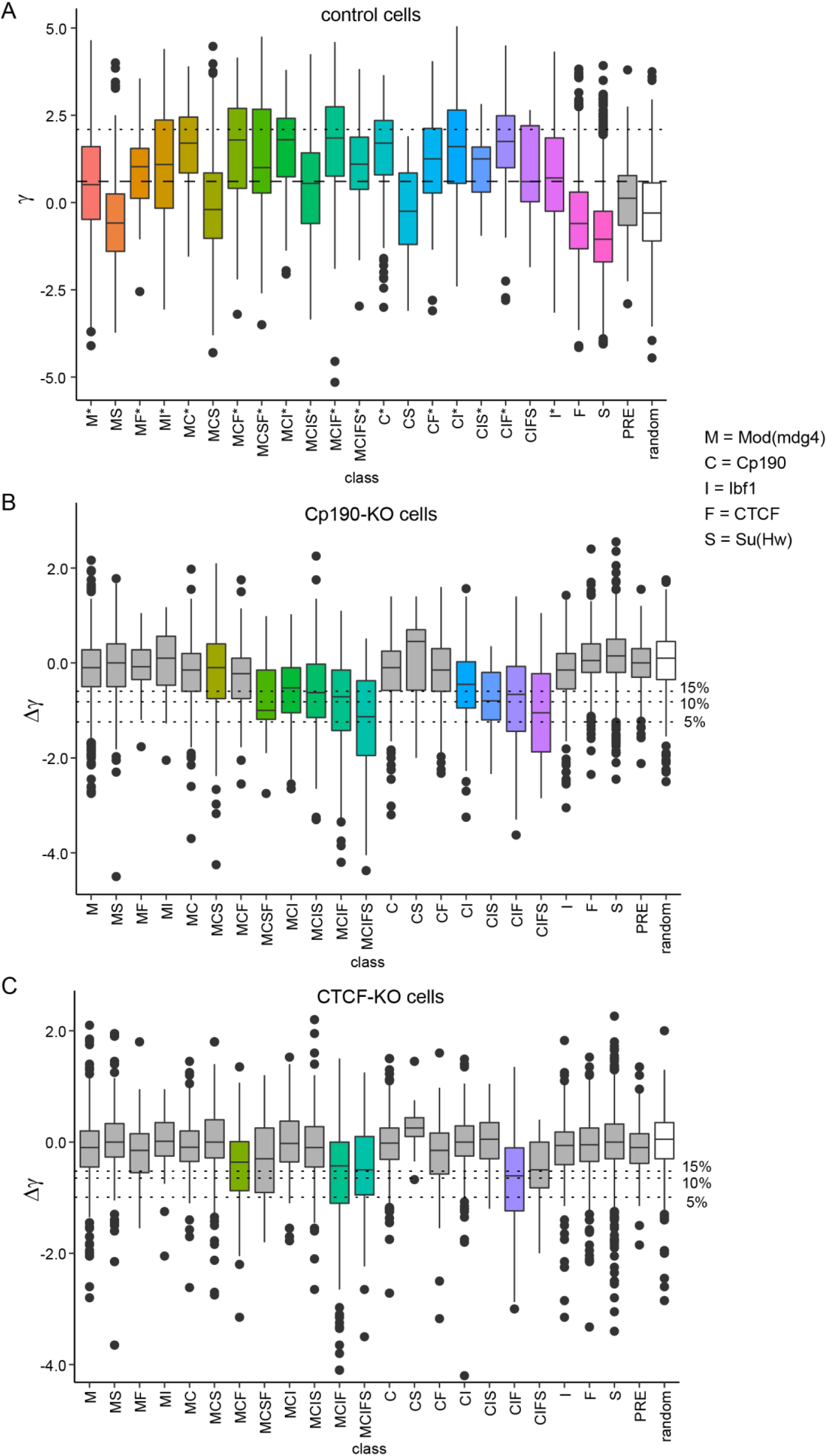
Specific combinations of insulator proteins limit chromatin contacts. **A.** Box-plots display distance scaling factors (γ) at different classes of insulator protein binding sites (coloured boxes), Polycomb Response Elements (PREs, grey box) and control regions (random, white box). Here and in all subsequent figures, the box-plots indicate the median and span interquartile range with whiskers extending 1.5 times the range and outliers shown as black dots. Sites with γ above the top quartile in the control group (horizontal dashed line) are considered as limiting chromatin contact crossing. Classes for which the fraction of such sites is significantly greater than that in the control group (*p*-value<0.0001, one-tailed Fisher’s exact test) are marked with asterisks. Horizontal dotted line indicates the top 5% value for γ genome-wide used to define TAD borders by Sexton and colleagues (*9*). Knock-out of Cp190 (**B**) and CTCF (**C**) leads to systematic reduction of distance scaling factors (negative Δγ = γ_KO_ - γ_control_) at some classes of insulator protein binding sites. The control group was used to define False Discovery Rates (% FDR, horizontal dotted lines). Classes for which the fraction of sites with Δγ below the 10% FDR cut-off significantly exceeds that in the control group (*p*-value<0.0001, one-tailed Fisher’s exact test) are marked with colour.

The 95^th^ percentile for γ genome-wide has been used as a threshold to define TAD borders (*9*). By this criterion, there are 1008 TAD borders in the control cells consistently identified in both replicate experiments (Figure S6). Of those, 913 correspond to one of the insulator protein bound regions, which means the majority (90.6%) of the TAD borders bind one or more known insulator proteins. In contrast, only 14.8% of all insulator protein bound regions are identified as TAD borders. Although, many of the insulator protein bound regions not identified as TAD borders are harder for chromatin contacts to cross compared to the control group (Figure 4A). These observations illustrate that it is difficult to pick a γ-based threshold that will reliably single out chromatin insulator elements.

Focusing on increase in chromatin contacts across insulator protein binding regions in Cp190-KO or CTCF-KO cells may be a better option. The change is described by the difference between distance-scaling factors calculated from the Hi-C measurements in mutant and control cells (Δγ=γ_KO_-γ_control_). As expected, the median Δγ for the control (random) regions is close to zero for both Cp190- and CTCF-KO (Figures 4B-C). Although, at some of the regions, Δγ deviates due to technical variability of Hi-C as well as inherent variability of cultured cell lines. In contrast, several classes of insulator protein bound regions show systematic increase in chromatin contact crossing (negative Δγ) upon Cp190- or CTCF knock-out (Figures 4B-C; *p*-value <0.0001, one-sided Fisher exact test). Importantly, the regions that, in the wild-type cells, bind Cp190 but no CTCF (e.g. MCS, MCI, MCIS and CIS) become systematically easier to cross (negative Δγ) only in the Cp190-KO (Figure 4B) but not in the CTCF-KO cells (Figures 4C). This indicates that our assay is specific.

To single out putative chromatin insulator elements, we followed a two-step algorithm. First, using the distribution of Δγ values for the random control regions we defined 5%, 10%, and 15% False Discovery Rate (FDR) thresholds. Second, using these thresholds we selected all regions with Δγ ≤ FDR from classes that have a significantly higher fraction of regions with increased chromatin contact crossing in the corresponding mutant cells (marked with colour on Figures 4B-C, see *materials and methods* for calculations). This way, we detected 745 putative insulator elements that require Cp190 or CTCF or both at 15% FDR (632 at 10% FDR; 401 at 5% FDR). For additional statistics and the list of elements, see Figure 3 and Table S3. This catalogue includes *Fub*, *62D, 1A2*, *SF1* and *Homie* insulator elements identified by genetic assays (*19, 22, 39, 44, 55, 59-61*). Approximately one third of the insulators from our catalogue (from 28.99% of the elements defined at 15% FDR to 30.67% - defined at 5% FDR) reside within 2kb from their nearest TAD border.

### Other factors impair chromatin contact crossing

Most classes of insulator protein bound regions, which show an increase in chromatin contact crossing in the mutant cells (Δγ < 0), are hard to cross (have high γ) in the control cells (Figure 4). However, the inverse is not true. For example, regions of the C, MC and CF classes bind Cp190, have high γ but show no increase in contact crossing upon Cp190 knock-out (Figure 4). Clearly, at these regions, other features can substitute for Cp190 or are the primary cause for the reduced chromatin contact crossing. What could these features be? BEAF-32 protein was implicated in the function of the *scs’* insulator element (*62*) but also in transcriptional activation (*63*). Many of the BEAF-32 binding sites overlap Cp190 bound regions ((*37*), Figure S7C). Nevertheless, when we compare the class C regions that bind BEAF-32 to those that do not bind the protein, it is evident that BEAF-32 is not the primary cause of high γ at these regions (Figures 5A-B).

**Figure 5.**
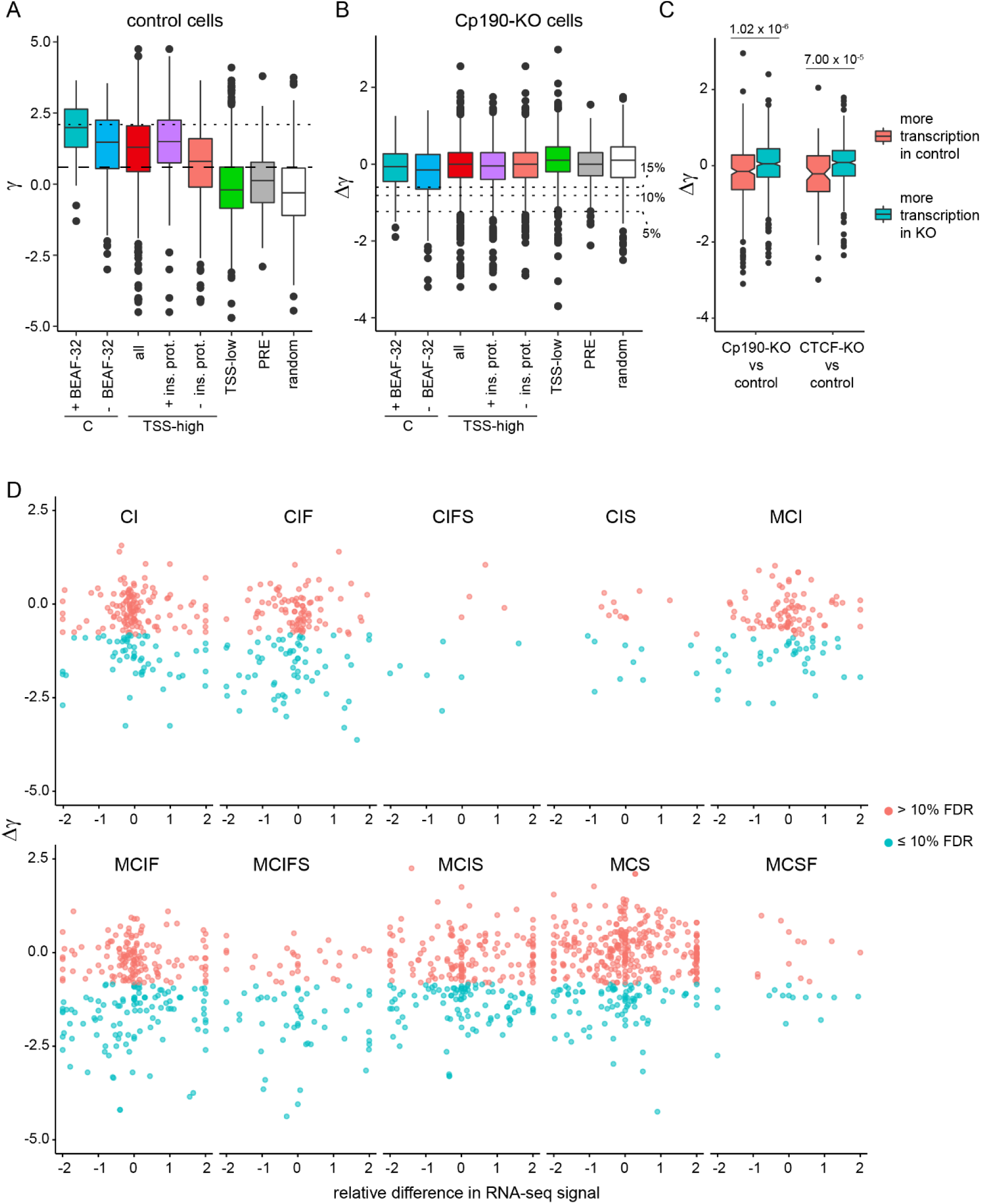
TSS of transcriptionally active genes impair chromatin contact crossing. **A.** Box-plots display distance scaling factors (γ) in control cells at class C regions either co-bound or not bound by BEAF-32. Also shown are γ of TSS grouped by transcriptional activity of the corresponding gene and the binding of insulator proteins mapped in this study. Distance scaling factors of PREs and randomly selected regions with no appreciable ChIP-seq signal for any of the insulator proteins are shown for comparison. Sites with γ above the top quartile in the random control group (horizontal dashed line) are considered as limiting chromatin contact crossing. Horizontal dotted line indicates the top 5% value for γ genome-wide. **B.** Cp190 knock-out leads to no systematic change in distance scaling factors (Δγ = γ_KO_ -γ_control_) at any of the classes of regions. Horizontal dotted lines indicate FDRs used to define insulator elements in Figure 4. **C.** Box-plots of Δγ for TSSs of genes whose transcription differs (|log_2_fold change| > 2) between Cp190-KO or CTCF-KO and control cells. Note significant difference between Δγ for TSS with higher transcription in the mutant cells compared to that of the genes with higher transcription in the control cells (*p*-values from Wilcoxon rank sum test are shown above). **D.** Scatter-plots compare the changes in distance scaling factors (Δγ ≤ 10% FDR coloured in cyan; Δγ > 10% FDR coloured in pink) at insulator protein binding sites of various classes to relative differences in RNA-seq signals of genes closest to these sites after Cp190 knock-out.

Further inspection of the C class indicates that most of these regions reside close to Transcription Start Sites (TSS) with median distance of just 226bp (Figure S7A). Sequencing of the total RNA indicates that most of the corresponding TSS belong to transcriptionally active genes (Figure S7B). Conceivably, proteins associated with transcription or spatial interactions between transcriptionally active genes restrict chromatin contact crossing regardless of Cp190 binding. Indeed, the Hi-C bins encompassing “highly transcriptionally active” TSS (top quartile of the RNA-seq signals) have γ comparable to that of the C class (Figure 5A). Many insulator protein bound regions are located close to transcriptionally active TSS (Figures S7A-B) possibly contributing to the high γ of the latter. Nevertheless, TSS of highly transcribed genes that have no Mod(mdg4), Cp190, Ibf1, CTCF, Su(Hw) or BEAF-32 bound within 1kb distance are still hard for chromatin contacts to cross (Figure 5A). Supporting this notion, chromatin contacts across a TSS whose transcription differs between Cp190-KO (or CTCF-KO) and control are harder to establish in cells where it is more transcriptionally active (Figure 5C).

Two conclusions follow from the observations above. First, transcriptionally active genes hinder chromatin contacts regardless of their association with Cp190 or any other insulator protein tested here. This may be due to binding of other, possibly undiscovered, insulator proteins or spatial segregation of transcriptionally active genes. Second, a previously recognised link between Cp190 and TAD boundaries (*9, 64*) should be interpreted with caution as large fraction of Cp190-bound regions resides next to TSS of transcriptionally active genes.

### Changes in chromatin contacts across insulator elements are not linked to altered transcription

Since transcriptionally active genes hinder the chromatin contact crossing, we wondered to what extent the increased contacts across insulator protein bound sites reflect changes in transcription of the nearby genes. Using the DESeq2 algorithm (*65*), we identified 831 genes differentially transcribed in Cp190-KO cells compared to control cells, 475 genes differentially transcribed in CTCF-KO cells compared to control and 691 genes differentially transcribed between Cp190-KO and CTCF-KO cells. Principle component analysis (PCA) indicates that the three cell lines have distinct changes in gene transcription with Cp190-KO cells being slightly more different from either CTCF-KO or control cells (Figure S8A).

Arguing against the link between topological changes at insulator sites and transcription, we found no coherent transcriptional changes in Cp190- and CTCF-KO cells around insulator elements impaired in both cell lines. As illustrated by clustering of the top twenty most differentially transcribed genes (Figure S8B), some of the observed variability is due to the stochastic nature of cell line derivation as well as distinct genetic backgrounds of parental fly lines. For example, note that *yellow* (*y*), the gene most “upregulated” in CTCF-KO cells (Figure S8B), comes from the transgene inserted in the *CTCF^y+1^* allele (*39, 42*).

More importantly, we see no correlation between the changes of distance-scaling factor (Δγ) at insulator protein binding sites and the transcription from the closest TSSs (Figures 5D, S9). Taken together these observations argue that the increased contacts across insulator protein binding sites are not linked to changes in transcriptional activity of the nearby genes caused by the ablation of Cp190 or CTCF. Instead, they are the direct consequence of the disrupted function of the underlying insulator elements.

### Chromatin insulator mode of action

The extensive catalogue of putative insulators may yield mechanistic insights in their function. As the first step towards this aim, we asked: from what distance could the contacts across insulator elements be blocked? To this effect, we calculated the number of contacts between pairs of Hi-C bins around each putative insulator starting from the two bins immediately adjacent to the insulator and followed by pairs at progressively larger distances (Figure 6A). We then subtracted the values for corresponding bin pairs calculated for mutant and control cells.

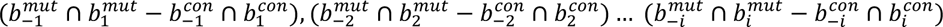

**Figure 6.**
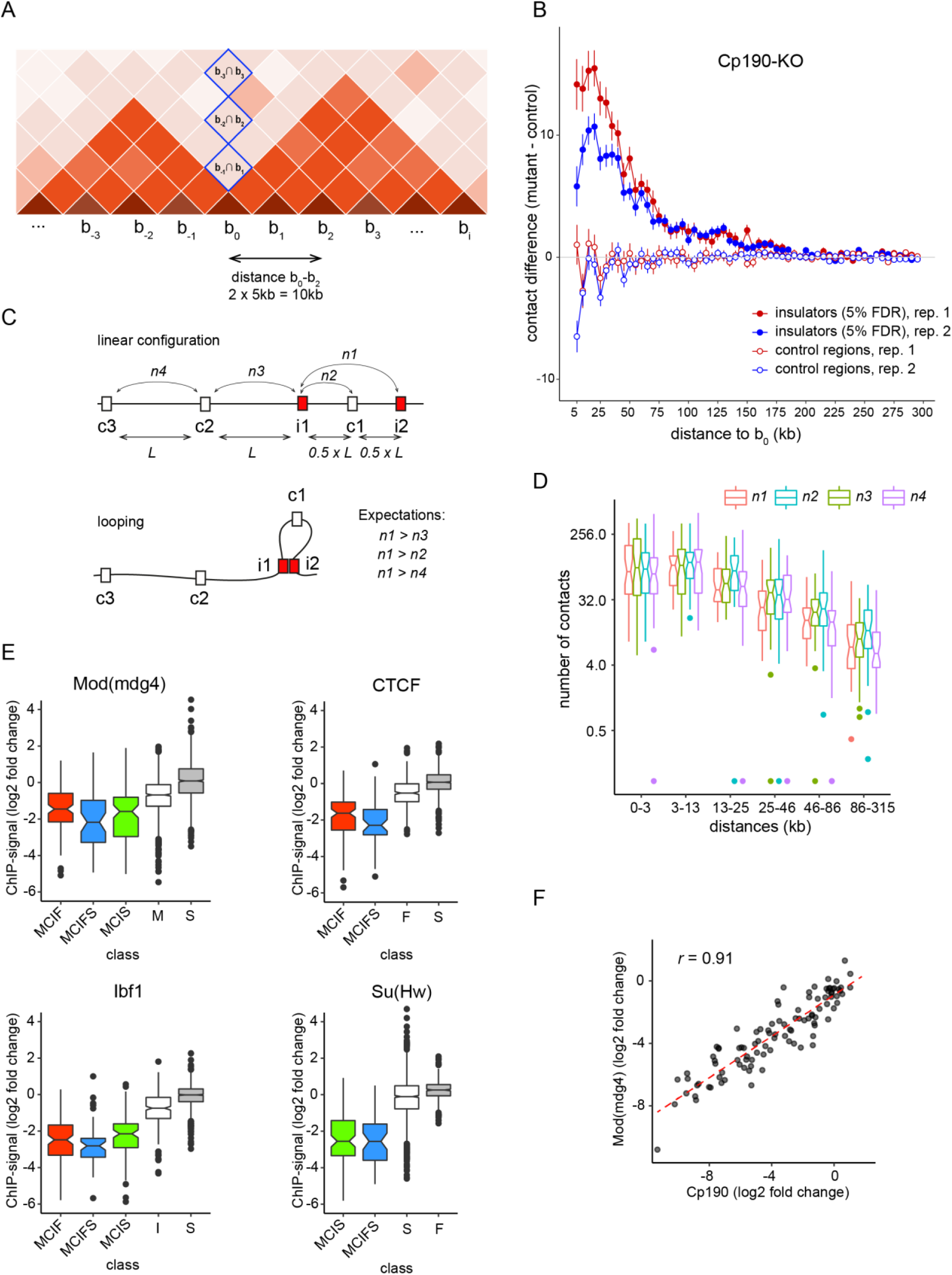
Action range and looping test. **A.** Schematic representation of a contact matrix segment to illustrate the metrics used in (B). The higher red colour intensity indicates more contacts. To estimate the difference in contacts across a region (e.g. chromatin insulator), we calculated the number of contacts between pairs of Hi-C bins around this region (bin b_0_) starting from the two bins immediately adjacent to the insulator (b_-1_ ∩ b_1_) and followed by pairs at progressively larger distances (b_-2_ ∩ b_2_, etc.). We then subtracted the values for corresponding bin pairs calculated for mutant and control cells to yield the contact crossing difference curves. Blue rectangles mark the contact matrix elements included in the curve. **B.** Average contact crossing difference curves for Cp190-dependent insulator elements (filled circles) and randomly selected control regions that do not bind any insulator proteins (empty circles) determined from two replicate Hi-C experiments (indicated with red and blue colours). Note that at close distances (5-10kb) the estimate of chromatin contact frequency from proximity ligation (the underlying principle of Hi-C method) becomes less reliable because, most of the time, chromatin fragments are sufficiently close to each other and the likelihood of successful ligation is more dependent on random chance. **C.** Schematic illustration of the insulator looping test. For each pair of the closest insulator elements (i1 and i2) located at a distance *L*, the number of contacts between the insulators (*n1*) is compared to that (*n2*) between the insulator i1 and the region c1 located half way between insulators i1 and i2. *n1* is also compared to the number of contacts (*n3*) between the insulator i1 and the region c2 or to the number of contacts (*n4*) between the two control regions c2 and c3, both separated by the distance *L*. A preferential looping interaction between the insulators i1 and i2 is expected to cause more frequent contacts between the two compared to all other pairs. **D.** Pairs of closest Hi-C bins containing insulator elements (defined at 15% FDR) were split in groups of equal sizes depending their separation in the linear genome. The number of contacts between the paired insulator bins (*n1*) was plotted (red box-plots) alongside the number of contacts between corresponding pairs of control bins (blue (*n2*), green (*n3*) and purple (*n4*) boxplots). Notches mark 95% confidence intervals of the medians. **E.** Box-plots of log_2_(Cp190-KO/control) changes in ChIP-seq signals at MCIF (red), MCIFS (blue) and MCIS (green) insulator sites. These are compared to standalone sites for the protein of interest (white) and the background noise at sites not significantly enriched by this protein (grey). No overlap between the box-plot notches indicates that their medians are significantly different. **F.** Scatter-plot of log_2_(CTCF-KO/control) changes of ChIP-seq signals for Cp190 and Mod(mdg4) at MCIF insulators. Dashed red line shows the linear regression fit.

The resulted contact crossing difference values were averaged for all putative insulator elements to yield the cumulative insulator contact crossing difference curves (Figures 6B, S10). To control for potential sampling and normalization biases we applied the same procedure to a set of randomly chosen regions that do not bind any of the insulator proteins. As expected, the cumulative contact crossing difference curves for control regions fluctuate around zero (Figures 6B, S10). In contrast, the curves for the putative insulator elements are positive up to the distances of ∼150kb (Figures 6B, S10). This argues that an average *Drosophila* insulator element can interfere with contacts between chromosomal sites that are up to 300kb apart.

How *Drosophila* insulator elements interfere with chromatin contacts is not well understood. The most popular hypothesis suggests that fly insulators physically interact with each other and form chromatin loops which, in turn, compete with chromatin contacts between chromosomal elements outside the loops. With an extensive catalogue of insulator elements at hand, we sought to evaluate this hypothesis using a “looping test” illustrated on Figure 6C. For each pair of the closest insulator elements from our catalogue, we calculated the number of contacts between Hi-C bins containing these insulators and between three matched control bin pairs. The first control pair consisted of one of the insulator bins (i1) and the control bin (c1) half way towards the second insulator (i2). The second control pair consisted of the insulator bin (i1) and the control bin (c2) located at the same distance as the two insulator bins. Finally, the third control pair included control bins c2 and c3 located at the same distance as the insulator bins. If insulator elements tend to interact with each other and form loops, we expect the number of contacts between the closest insulator pairs to be greater than that between the control bin pairs (*n1* > *n4*; *n1* > *n3*; *n1* > *n2*). The box plots of contact numbers (Figure 6D) indicate that, regardless of genomic distances between the closest insulators, this is not the case. To summarise, the results of our test provide no support for the model where *Drosophila* insulator elements block chromatin contacts by forming chromatin loops.

One may expect that sites bound by multiple insulator proteins would impair chromatin contacts even when Cp190 or CTCF are missing because the other proteins would compensate for their loss. Our experiments indicate the opposite (note the low Δγ for MCIF, MCIFS, MCIS sites, Figures 4B-C). Two not mutually exclusive explanations may account for this. First, the ablation of Cp190 or CTCF may lead to the loss of other insulator proteins from these sites. Second, at sites co-bound by multiple insulator proteins, simultaneous presence of all proteins may be required to limit the chromatin contact crossing. To evaluate these possibilities, we compared the ChIP-seq signals for Mod(mdg4), Ibf, CTCF and Su(Hw) at MCIF, MCIFS, MCIS insulators between the Cp190-KO and control cells. For all proteins, the immunoprecipitation of these regions from the Cp190-KO chromatin is significantly reduced compared to that of the controls (Figure 6E). This indicates that Mod(mdg4), Ibf, CTCF or Su(Hw) require Cp190 for efficient binding to MCIF, MCIFS and MCIS sites. The CTCF knock-out also affects Cp190 and Mod(mdg4) binding to the MCIF insulators to an extent that varies between individual sites. Further strengthening the Cp190 dependence argument, the reduction of Cp190 and Mod(Mdg4) ChIP-seq signals upon CTCF ablation is highly correlated (Figure 6F). To summarize, it appears that the loss of Cp190, by mutation or due to impaired tethering by CTCF, impairs the binding of companion insulator proteins, which explains why these proteins do not compensate for Cp190 loss.

## Discussion

Three main conclusions follow from our study. First, the *Drosophila melanogaster* genome contains hundreds of Cp190- and/or CTCF-dependent chromatin insulators. While they appear to play relatively minor role in shaping the overall chromosome folding patterns, they have distinct impact on chromatin contacts at many specific loci. Second, we find that TSS of transcriptionally active genes are generally hard for chromatin contacts to cross regardless of their association with Cp190 or any other insulator protein that we tested. Since many Cp190-bound regions reside next to TSS of transcriptionally active genes, a previously recognised link between Cp190 and TAD boundaries should be interpreted with caution. Third, the expanded catalogue of insulator elements is instrumental to advance our understanding of how these elements affect chromatin contacts. For example, we found no evidence that Cp190- or CTCF-dependent insulators preferentially interact with each other. This argues that new models, which do not invoke chromatin loops formed by interacting insulator elements, are required to explain the mechanism of insulator action. Broadly similar conclusions were reached by the study of Kaushal and co-authors (*66*) who reported the analysis of genome folding in *Drosophila* embryos deficient for Cp190 and CTCF while this work being prepared for publication.

Discovered some 30 year ago, chromatin insulators were hailed as major players organizing the *Drosophila* genome into topologically independent regulatory units. This outlook started to fade as we learned more about the architecture of the fly chromatin. It became apparent that transcriptional activity is a better predictor of the overall *Drosophila* TAD organization than genomic distribution of insulator proteins (*67*) and that partial depletion of insulator proteins by RNAi causes only limited changes in patterns of posttranslational histone modifications or gene transcription (*37, 68*). It was then hypothesized that topological partitioning of the *Drosophila* genome is driven primarily by interactions between transcriptionally active genes (*35, 67*) and that, due to its much more compact genome, flies may not require additional mechanisms to regulate genome architecture (*35, 69*). Consistent with this view and in contrast to vertebrates, there is no evidence that *Drosophila* TADs arise from blocking cohesin-mediated chromatin loop extrusion.

Our work reconciles the two assessments. On one hand, our Hi-C analyses indicate that complete ablation of Cp190 or CTCF does not grossly disrupt the genome folding. On the other hand, we show that *Drosophila* genome contains over 700 putative insulator elements, which restrict chromatin contact crossing and can interfere with contacts between chromosomal sites up to 300kb apart. As we know from prior genetic analyses some of these insulators, e.g. *Fub*, are essential for correct regulation of developmental genes. These insulator elements restrict chromatin contacts regardless of transcriptional activity within the neighbouring chromatin. As exemplified by *Fub*, they can impair chromatin contacts within a locus repressed by the Polycomb mechanisms. While transcription related mechanisms appear to define the major contact patterns within the *Drosophila* genome, insulators have widespread but shorter-range impact.

As clear from observations presented here and those by Kaushal and co-authors (*66*), at many sites co-bound by CTCF and Cp190, the former contributes to Cp190 recruitment to chromatin. Yet, ChIP-seq analysis of sites bound by multiple insulator proteins (e.g. of MCIF, MCIFS and MCIS classes) indicates that, at these locations, Cp190 is required for efficient binding of other insulator proteins. Surprisingly, those include the sequence specific DNA-binding proteins CTCF and Su(Hw). Three not mutually exclusive possibilities can account for this observation. First, it is possible that Zn-finger domains of Cp190 increase the overall affinity of the multi-protein insulator complex via sequence-unspecific binding to DNA (*70, 71*). Second, Cp190 forms dimers (*72, 73*), with each molecule capable of interacting with its own sequence-specific DNA binding partner e.g. Su(Hw) and CTCF or CTCF and Ibf etc. This, in turn, would allow for Cp190-dependent cooperative binding of the whole complex to DNA. Third, Cp190 was reported to interact with Nurf301, the core component of the NURF chromatin remodelling complex (*74*). The NURF complex slides nucleosomes, which may enable Cp190-associated proteins to bind their cognate sequence motifs more efficiently. Additional experiments will be required to discriminate between these possibilities.

It has been widely assumed that *Drosophila* insulator elements restrict chromatin contacts by interacting with each other and forming chromatin loops. This model is appealing, as it would explain several phenomena observed in transgenic experiments, for example, the “insulator bypass” (*28, 29*), the insulator-mediated long range transvection (*21, 75*) and the trans-interactions between epigenetic regulatory elements (*76*). Of note, in the latter two cases, the interactions involve insulators and transcription. Since some of the insulator proteins can homodimerize, the cognate DNA elements could interact and form metastable loops leading to “bypass” when insulators are located nearby within a transgenic construct. Similarly, otherwise weak insulator-mediated interactions may be strengthened by mechanisms, which drive interactions between transcriptionally active genes, together enabling long-range transvection. Regardless, our observations argue that the insulator interactions do not happen or do not persist long enough to be distinguished from the genomic average, when these elements reside in their endogenous locations. It is, therefore, hard to imagine that such transient or infrequent interactions would be sufficient for *Drosophila* insulator elements to block chromatin contacts by forming chromatin loops. This conclusion agrees with results of recent Micro-C experiments from Levine’s laboratory who reported that looping interactions within the *Antennapedia* gene complex do not involve any of the known or putative insulator elements (*77*). Using the HiCCUPS algorithm (*78*), Chathoth and co-authors reported several hundred loops formed by regions co-bound by Cp190, BEAF-32 and Chromator (*68*). These sites are predominantly TSS of active genes. The majority of such sites were not classified as insulator elements in our screen because their impact on chromatin contact crossing does not require Cp190 or CTCF. While we cannot exclude that certain active TSS contain elements that impair chromatin contacts by forming loops, additional experiments would be needed to uncouple this effect from generic interaction between transcriptionally active genes.

To conclude, the extended catalogue of insulator elements uncovered in our study will provide a significant new resource to study the regulation of specific *Drosophila* genes as well as general mechanisms that shape the folding of eukaryotic genomes.

## Materials and methods

### Derivation and culture of Cp190-KO and CTCF-KO cell lines

The *CTCF^y+1^* and *Cp190^3^* fly strains (*42, 43*) were used to derive the corresponding mutant cell lines following the procedure of Simcox et al. (40) with modifications described in (*46*). Cells were cultured at 25^0^C in Schneider’s media (Lonza), supplemented with 10% heat-inactivated fetal bovine serum (Sigma), 0.1mg/ml streptomycin and 0.1u/ml penicillin (Gibco) under sterile conditions.

### Hi-C, library preparation and sequencing

Hi-C was performed as described in (*9*). Briefly, 2×10^7^ cells were crosslinked by incubating in fix buffer (2% formaldehyde, 15mM HEPES pH 7.6, 60mM KCl, 15mM NaCl, 4mM MgCl2, 0.1% Triton-X100, 0.5mM DTT, protease inhibitor cocktail (Roche)) for a total of 10 min at 25°C, 750 rpm on a shaker. After quenching with 5ml of 2M glycine, permeabilized cells were collected by centrifugation for 5 min at 4500 g, 4°C, then washed once with 5ml fix buffer (without formaldehyde) and once with 1.25 x NEB3 buffer (New England Biolabs), with centrifugation for 5 min at 4500g, 4°C each time. Permeabilized cells were resuspended in 300µl 1.25 x DpnII buffer (New England Biolabs) plus 0.3% SDS and incubated for 1 hr at 37°C, 1000 rpm on a shaker. Triton-X100 was added to a final concentration of 2% and the permeabilized cells were incubated for a further 1 hr at 37°C, 1000 rpm, before overnight treatment with 1500U of DpnII (New England Biolabs) at 37°C, 1000 rpm. The restriction enzyme was inactivated by incubation for 20 min at 65°C, 1000 rpm with SDS at a final concentration of 1.3%, before dilution of the lysate in 10ml of 1x T4 DNA ligase buffer (New England Biolabs) plus 1% Triton-X100 and incubation for 1 hr at 37°C, 750 rpm. The released chromatin was ligated for 4 hr at 25°C, 750 rpm with 40,000 U T4 DNA ligase (New England Biolabs). Then crosslinks were reversed overnight at 65°C, 750 rpm in the presence of 150µg/ml proteinase K. The resulting 3C DNA was purified by 1 hr treatment with 40µg/ml RNase A at 37°C, 750 rpm, phenol extraction, phenol/chloroform extraction and ethanol precipitation. The DNA was quantified with the Qubit dsDNA assay (Invitrogen). 5µg aliquots of 3C DNA were sonicated with Bioruptor (Diagenode) in 50 µl volumes in sonication buffer (50 mM Tris-HCl, pH 8, 10 mM EDTA, 1% SDS) to obtain a fragment range between 500-1500bp. The sonicated 3C DNA was then purified by phenol/chloroform extraction and ethanol precipitation and quantified with the Qubit dsDNA assay (Invitrogen). Libraries for paired-end sequencing were made from 500 ng aliquots of sonicated 3C DNA using Illumina reagents and protocols, with size selection for products of ∼800 bp. The libraries were sequenced on a HiSeq 2000 instrument (Illumina), following the manufacturer’s protocol.

### ChIP library preparation and sequencing

ChIP and qPCR analysis were done as described in (*46*) except for chromatin was sonicated in 4ml of 10mM Tris-HCl pH8.0, 1mM EDTA pH8.0 with Branson D450 sonicator for 45 minutes (forty five cycles of 20 sec ON – 40 sec OFF, amplitude 40%) and adjusted to 5ml in RIPA buffer (10mM Tris-HCl pH8.0, 1mM EDTA pH8.0, 1% Triton-X100, 0.1% SDS, 0.1% DOC, 0.14mM NaCl). Isolated ChIP DNA was re-suspended in 40µl of DNAse free water and used for ChIP-seq library preparation. 4µl of precipitated DNA were diluted 10 fold and used for qPCR analysis to check the specificity of ChIP reactions. The antibodies used are listed in Table S4 and the nucleotide sequences of qPCR primers in Table S5. For ChIP-seq library preparation 2ng of immunoprecipitated DNA were processed using NEBNext Ultra II DNA Library preparation Kit for Illumina (cat# E7645) and index oligonucleotides from NEBNext Multiplex Oligos for Illumina (cat# E7335). Fragments of average size of 180bp were selected with SPRIselect® Reagent Kit (Beckman Coulter, Inc. #B23317), amplified for 15 cycles, pooled and sequenced (10 libraries per one flow cell) at SciLifeLab national sequencing facility (Stockholm branch) with HiSeqX instrument (HiSeq Control Software 2.2.58/RTA 1.18.64) and 1×51 setup using ’HiSeq SBS Kit v4’ chemistry. The Bcl to FastQ conversion was performed using None from the CASAVA software suite. The sequence read quality was reported in Sanger/phred33/Illumina 1.8+ scale.

### Strand-specific total RNA library preparation and sequencing

Total RNA was isolated from cultured cells using TRI Reagent according to the manufacturer instructions (Sigma-Aldrich #T9424). 200ng of total RNA were used for indexed library preparation using Ovation RNA-Seq Systems 1–16 for Model Organisms Kit (#0350 DROSOPHILA, Nugen Technology). Briefly, total RNA was treated with DNase I and reverse transcribed using random oligonucleotide primers. The resulted cDNA was fragmented to 200bp with Covaris E220 focused-ultrasonicator using micro-tubes (AFA Fiber Crimp Cap), the cDNA fragments were end repaired, ligated to oligonucleotide adaptors, strand separated and subjected to 18 cycles of PCR amplification. After purification the libraries were pooled and sequenced at SciLifeLab national sequencing facility (Stockholm branch) using one flow cell of HiSeq2500 (HiSeq Control Software 2.2.58/RTA 1.18.64) with a 2×126 setup and ’HiSeq SBS Kit v4’ chemistry. The Bcl to FastQ conversion was performed using None from the CASAVA software suite. The sequence read quality was reported in Sanger/phred33/Illumina 1.8+ scale.

### Hi-C analysis

#### Primary data processing and normalization

Sequence read mapping to the *Drosophila melanogaster dm3* genome release (see statistics in Table S1), filtering, and normalization were performed as previously described (*9*). The resulted statistics on the number of observed contacts for each pair of restriction fragments and the number of expected contacts from a low-level probabilistic model, which considers local GC content and the Dpn II restriction fragment length (*79*) is reported in Table S2. Technically corrected contact matrices were generated by calculating ratios between the total observed reads and the expected reads based on the above model. Bin-contact pair maps were transformed to the *gcmap* format using *gcMapExplorer bc2cmap* with Iterative Correction (IC) *gcMapExplorer normIC* (*80*).

For pairwise comparisons of Hi-C contact matrices, Spearman’s rank correlation (ρ) and Pearson moment correlation (*r*) coefficients were calculated using *cor* function in R (https://www.R-project.org/). To avoid spurious experimental noise, the matrices were filtered to remove bins with observed contacts of less than two, contacts within individual bins (the diagonal of the contact matrix), and between bins separated by more than 1.6Mb were not considered. The similarity between Hi-C contact matrices of individual experiments was evaluated by hierarchical clustering as implemented in *hclust* function with distances between experiments calculated as 1-absolute value of correlation coefficient. The cluster stability was evaluated by performing four variations of the procedure i.e., using ρ or *r* and two agglomeration methods: complete and average (UPGMA). For bins equal or larger than 40kb, the clustering was robust to variations (Figures 1C, S4).

All comparisons between Hi-C, ChIP-seq and RNA-seq datasets were performed using datasets in *dm3* genomic coordinates. In addition, for visualisation and reporting purposes, positions of insulator regions and contact heat-maps were also transposed to the *dm*6 genome release coordinates using the UCSC *LiftOver* tool.

#### Data visualization

Contact matrices for replicate experiments at 5kb bin resolution were combined, converted to *gcmap* format, subjected to IC correction and displayed using *gcMapExplorer* browser (*80*).

#### *Distance-scaling factor* (γ) computation and assignment to specific regions

The *distance- scaling factor* (γ) was computed for each DpnII restriction fragment as described by (*79*). TAD borders were called from restriction fragment-level contact matrices of individual Hi-C experiments with control cells using the 95^th^ percentile for γ genome-wide as the threshold (*9*). The accuracy of TAD border positions was estimated by comparing the two replicate experiments (Figure S6) and was set to 2000bp. That is TAD borders identified in replicate experiments at a distance of 2000bp or less were considered identical. To assign γ values to insulator protein bound regions, coordinates of each region were compared to the coordinates of the DpnII restriction fragments and the overlapping fragments selected for further analysis. Insulator protein bound regions were segmented into 200bp sliding windows and coordinates of these windows compared to coordinates of the selected DpnII restriction fragments. *bedtools intersect -wa -wb* was used for all coordinate comparisons (*81*). From this, the weighted γ of each 200bp sliding window was calculated from γ of individual DpnII restriction fragments taking into account the degree to which the window overlapped these fragments. The highest value of all windows contained within an insulator protein bound region was taken to represent the region’s γ.

#### Definition of classes of insulator protein bound regions with systematic increase in chromatin contact crossing after Cp190- or CTCF-KO and calculation of False Discovery Rates (FDRs)

The systematic increase in contact crossing was defined as a negative shift in Δγ values, which was unlikely to happen by chance given the null distribution approximated by Δγ at 344 random sites not bound by any of the insulator protein profiled. To this end, the proportion of sites with bottom 10% of Δγ at a class of insulator protein regions was compared to that at the random sites using one-sided Fisher’s exact test. For classes of insulator protein regions that displayed systematic chromatin contact crossing upon Cp190- or CTCF-KO, Δγ values for bottom 5^th^, 10^th^ and 15^th^ percentiles among 344 random sites were used as corresponding FDR thresholds to call the regions significantly affected by knock-down (putative chromatin insulator regions).

#### Calculations of contact crossing difference curves

To account for sampling biases between Hi-C experiments, the contact matrices were further normalized using *multiHiCcompare* algorithm (*82*), which provides cyclic loess and fast loess methods adapted to jointly normalize the Hi-C data with more than two groups and multiple samples per group (Supplementary code file: *hic_normalize.R*). The normalized contact matrices were used to calculate numbers of contacts between pairs of Hi-C bins around selected central bins (b_0_) starting from the two bins immediately adjacent to b_0_ and followed by pairs at progressively larger distances of up to 60 bins (300kb = 60×5kb) (Supplementary code file: *gen_ContactCcrossingCurves.sh*). The values for corresponding bin pairs calculated for mutant and control cells were subtracted to yield the contact crossing difference curves.

#### Insulator looping test

Putative chromatin insulator sites were grouped by chromosome and sorted by their starting position to identify the closet pairs (i.e., *i1* and *i2*, as illustrated on Figure 6C). Each pair of the closest insulator elements (*i1* and *i2*) located at a distance *L*, were supplemented with three matching control bin pairs. The first control pair consisted of one of the insulator bins (*i1*) and the control bin (*c1*) located at a distance of *L/2* towards the second insulator bin (*i2*). The second control pair consisted of the insulator bin (*i1*) and the control bin (*c2*) located at the distance *L* to the left (upstream) of *i1*. Finally, the third control pair included control bins *c2* and *c3*, with the latter located at distance *L* to the left of *c2*. The number of contacts between bins of each pair were calculated using HiCmapTools (*83*) (Supplementary code file: *gen_InsulatorLoopingTest.sh*).

### ChIP-seq analysis

Reads were aligned to the *Drosophila melanogaster dm3* genome assembly using bowtie2 (*84*) and the following parameters *--phred33 -p 8*. Reads with MAPQ scores less than 30 were removed using samtools (*85*) *view* command and *-q 30*. Genomic read count profiles were computed from the filtered bowtie2 alignments using pyicos convert software (*86*) and the following parameters *-f sam -F bed_wig -x 180 -O*. The resulted read count profiles were normalized to the number of corresponding MAPQ 30 filtered sequencing reads using custom R script. Positions of ChIP-seq signal maxima within regions significantly enriched in both replicate experiments with the chromatin from control cells, were identified with MACS2 (v2.1.2) (*56*) *callpeak* command using the following parameters: *-f SAM -g dm --keep-dup all --fe-cutoff 8*. The resulted *bed* files were extended ± 300bp. Genomic positions of the regions above defined for each insulator binding protein were compared pairwise in order Mod(mdg4), Cp190, Ibf1, CTCF, Su(Hw). Thus, regions enriched by ChIP with antibodies against Mod(mdg4) were checked for overlap with regions enriched with antibodies against Cp190. The resulting three groups, i.e. bound by both Mod(mdg4) and Cp190, bound by just Mod(mdg4) or bound by just Cp190, were further compared to regions enriched by ChIP with antibodies against Ibf and so on. The resulted co-binding classes were designated with combinations of single letters representing individual insulator proteins bound.

To calculate ChIP-seq signal scores, read count profiles of individual insulator protein bound regions or Transcription Start Sites (TSS) were extracted from normalized genomic read count profiles of corresponding proteins using BEDTools (*81*) *intersect* function with parameters *-wa -wb*. The region-specific profiles were then used to calculate average read count per base pair ChIP-seq signal scores. TSS regions were defined by ± 1000bp extension of TSS positions obtained from UCSC Genome Browser (*87*).

### RNA-seq analysis

Paired-end reads were mapped to the *Drosophila melanogaster dm3* genome assembly using STAR_v2.6.1a (*88*) with default parameters. Unmapped or non-unique mapped reads were discarded (see statistics in Table S6). The gene transcription value was quantified as read count with *--quantMode GeneCounts*. Genes with transcription values within the top quartile were designated as transcriptionally active (TSS-high). Conversely, genes with transcription values within the bottom quartile were designated as transcriptionally inactive (TSS-low).

Differentially transcribed genes we identified using DESeq2 (*65*) with *log*2 fold change = 1 and Wald significance test *p*-value = 0.05 as thresholds. Principal Component Analysis (PCA) and identification of the top 20 most variable genes were performed after *regularized log* (rlog) transformation of transcription values.

## Data access

ChIP-seq, RNA-seq and Hi-C datasets generated in this study are available at Gene Expression Omnibus database under super series accession number GSE198763 with subseries: GSE198762, GSE198761, and GSE198760, respectively.

## Acknowledgements

We are grateful to Drs. R. Renkawitz, V. Corces, H. Saumweber and M.L. Espinas for generous gifts of antibodies. The work in YSC laboratory was supported in parts by grants from Swedish Research Council [2021-04435], Erik Philip-Sörensen Stiftelse, Carl Tryggers Stiftelse [CTS 12:434] and Knut and Alice Wallenberg Foundation [2014.0018, YBS co-PI]. Research in the J-M.C laboratory was supported by grants from the Taiwan Ministry of Science and Technology [108-2628-E-004-001-MY3] and would like to acknowledge the National Center for High-performance Computing (NCHC) for providing computational and storage resources. ChIP and RNA sequencing was performed by Science for Life Laboraroty at the SNP&SEQ Technology Platform in Uppsala. The facility is part of the National Genomics Infrastructure (NGI) Sweden and supported by the Swedish Research Council and the Knut and Alice Wallenberg Foundation.

---

**Figure S1.**
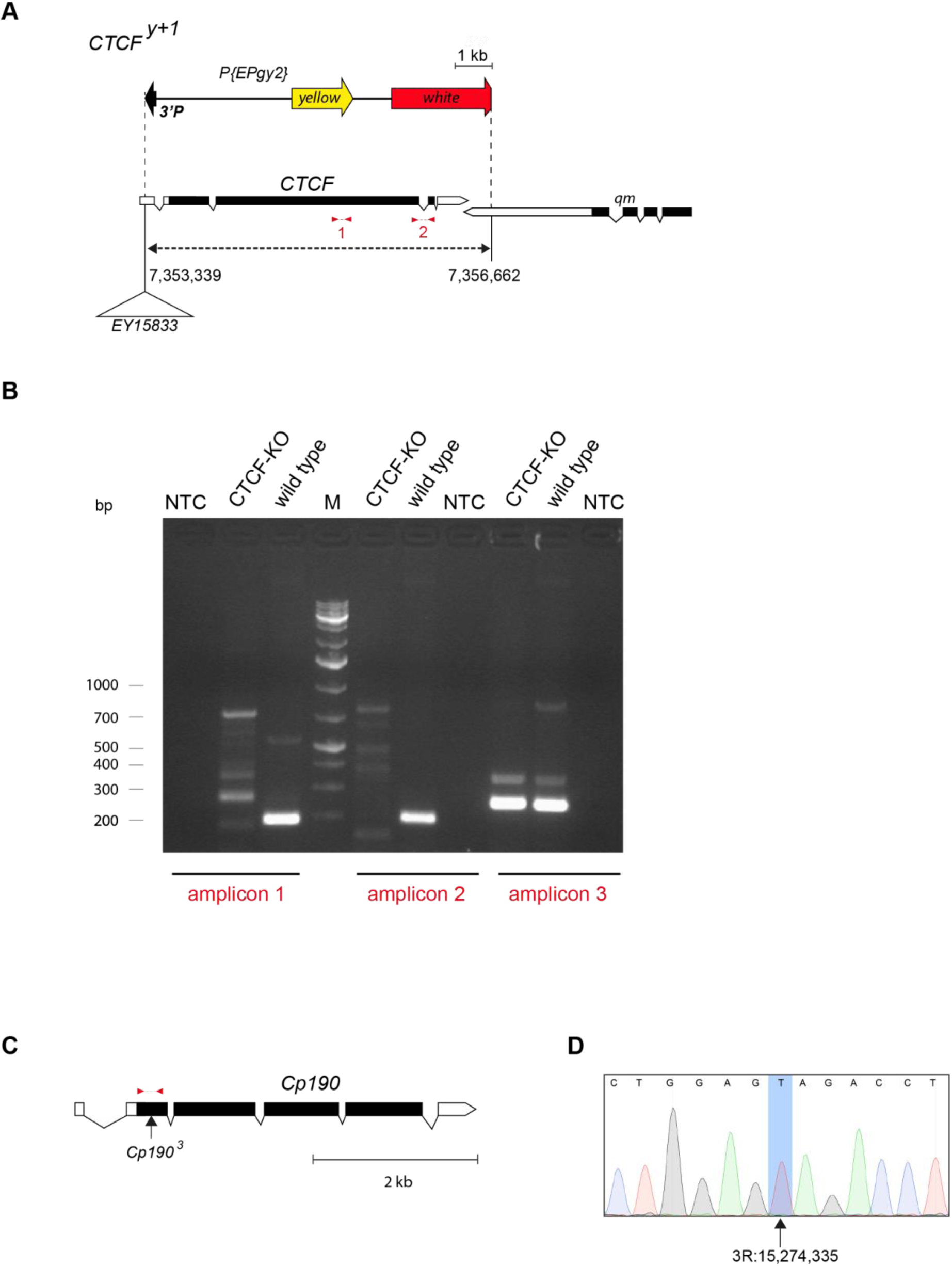
Characterization of CTCF 19.7-1c (CTCF-KO) and CP-R6 (Cp190-KO) cultured cell lines. **A.** Schematic representation of *CTCF ^y+1^* allele. The allele corresponds to 3.3kb deletion (dashed line) that removes most of the *CTCF* open reading frame and a small part of the 3’ UTR of the adjacent *qm* gene. Exons are shown as boxes, coding parts marked with back. Coordinates of deletion breakpoints are according to *Drosophila melanogaster* genome release BDGP R6/*dm6*, 2014. The *CTCF ^y+1^* deletion was generated by imprecise excision of *P[EPgy]CTCF^EY15833^* transposon (indicated as white triangle) and still contains some of its constituents including functional *yellow* gene and part of the *white* gene (shown above the molecular map of the *CTCF* locus). Note, that the schematic of *P[EPgy]CTCF^EY15833^*remnants is in different scale than that of the *CTCF* map. Positions of amplicons used to confirm the *CTCF ^y+1^* deletion in CTCF-KO cell lines are indicated with red lines. **B**. Genomic DNAs from CTCF 19.7-1c (CTCF-KO) and Ras 3 (wild type) cells were used for PCR with CTCFfw4 and CTCFrev3 primers (amplicon 1, expected product size - 189bp), CTCFfw5 and CTCFrev2 (amplicon 2, expected product size - 196bp) as well as BP3 and BP4 (amplicon 3, positive control outside the *CTCF* locus, expected product size – 258bp). As shown by electrophoresis of PCR products in 2% agarose gel, only reactions with genomic DNA from the wild-type cells yield single products corresponding to amplicons 1 and 2. This confirms that all CTCF-KO cells lack *CTCF* open reading frame. **C.** Molecular map of *Cp190* gene. Arrow indicates the position of C to T transition (Q61Stop) in the *Cp190^3^*allele. Red line indicates the location of the PCR product sequenced to genotype Cp190-KO cells. **D.** Genotyping of CP-R6 (Cp190-KO) cells. Genomic DNA from CP-R6 cells was amplified by PCR with CP190fw and CP190-Ncolrev primers, and the product of expected size – 382bp sequenced. The fragment of Sanger sequencing chromatogram illustrates that all CP-R6 chromosomes contain T instead C (blue shaded rectangle) at position 3R:15,274,335 (genome release BDGP R6/*dm6*, 2014).

**Figure S2.**
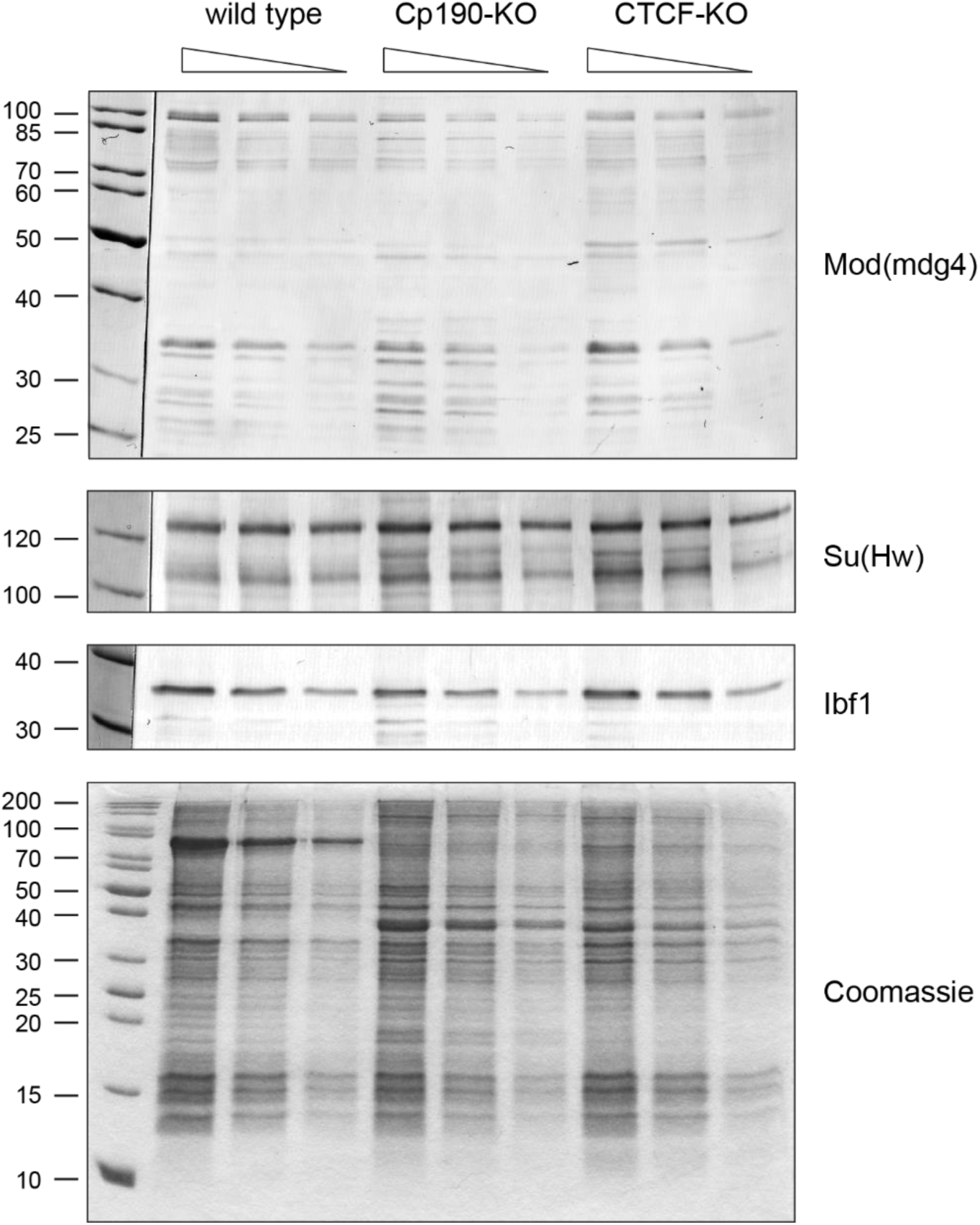
Ablation of Cp190 and CTCF does not affect the overall levels of other insulator proteins. Two-fold dilutions of total nuclear protein from control (Ras3), Cp190-KO (CP-R6) and CTCF-KO (CTCF 19.7-1c) cells were analysed by western-blot with antibodies against Mod(mdg4), Su(Hw) and Ibf1. Coomassie stained gel of corresponding total nuclear protein samples and western-blot with antibodies against Pc (Figure 1A) were used to control equal loading. Positions of molecular weight markers (in kDa) are indicated to the left.

**Figure S3.**
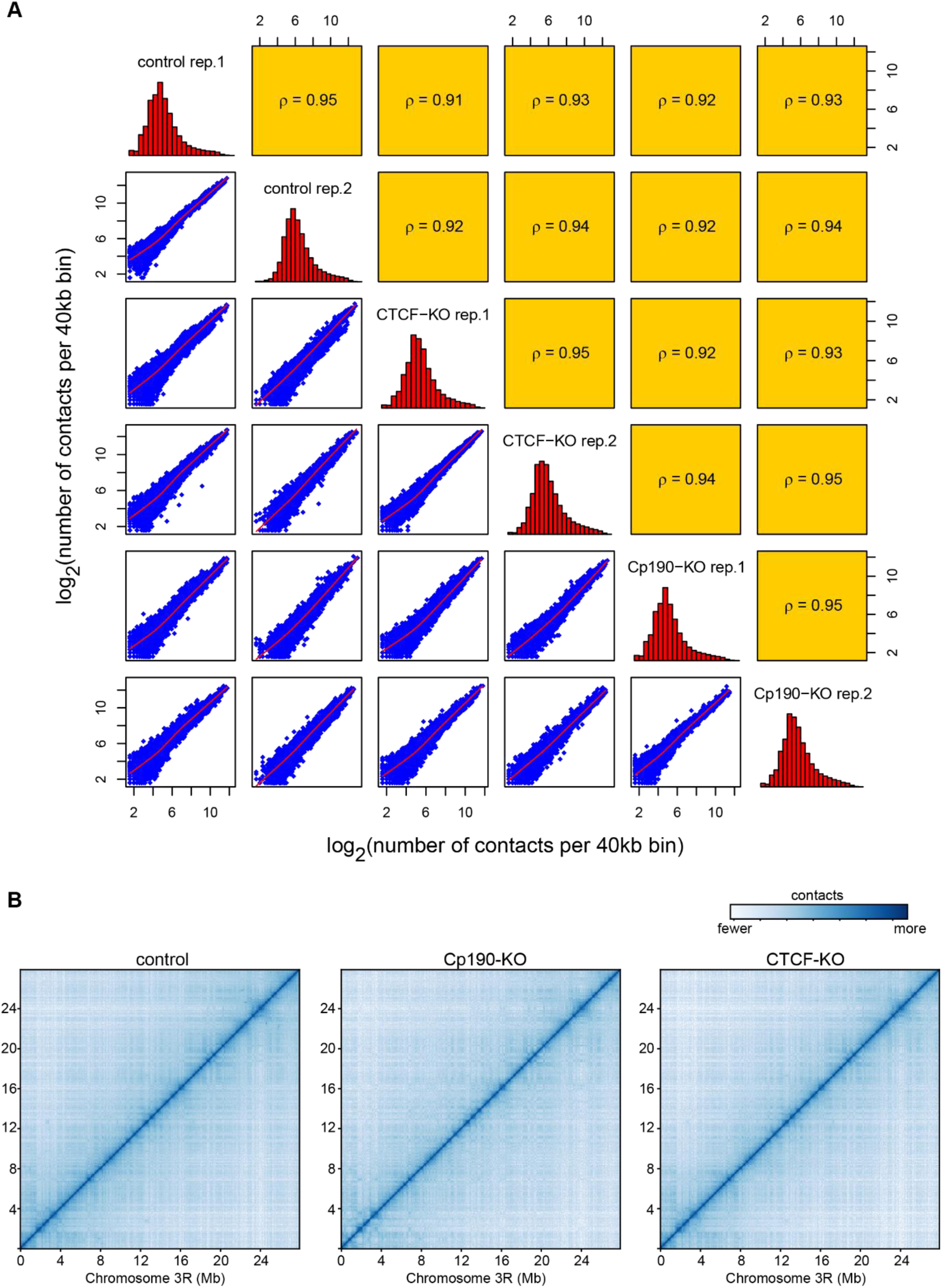
Overall chromatin contacts in the control, Cp190-OK and CTCF-KO cells are similar. **A.** Pairwise correlations between numbers of contacts for 40kb genomic segments (Hi-C bins) measured in individual experiments. Histograms along the diagonal show distributions of contact numbers for each experiment. Scatter plots below the diagonal illustrate the correspondence between contact numbers for individual bins (blue dots). Orange squares above the diagonal show corresponding Spearman’s rank correlation coefficients. To avoid spurious experimental noise, contacts within individual bins (the diagonal of contact matrix) as well as contacts between bins separated by more than 1.6Mb were not considered. **B.** Chromatin contacts over the right arm of *Drosophila melanogaster* chromosome 3 were counted at 40kb resolution, normalized by iterative correction (*89*) and plotted using *gcMapExplorer* software (*80*). To allow simultaneous visualization of the low- and high-frequency contacts, the heat map representation was scaled as logarithm of the map.

**Figure S4.**
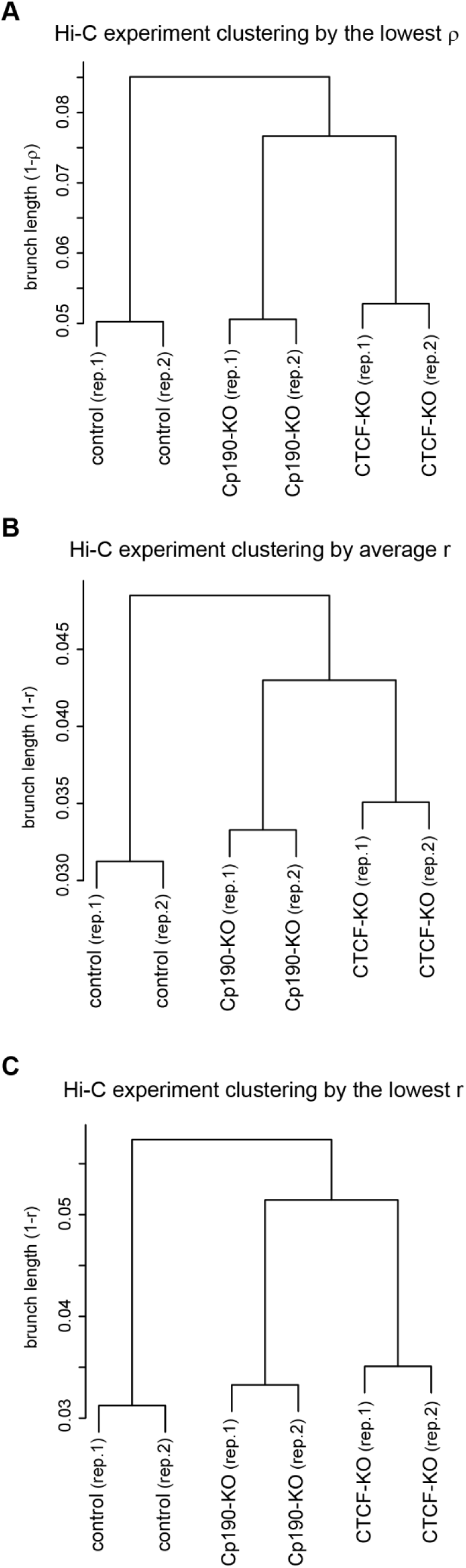
For bins equal or larger than 40kb hierarchical clustering of individual Hi-C experiments is robust to parameter changes. Chromatin contacts measured in individual Hi-C experiments were assigned to 40kb genomic segments (Hi-C bins) and clustered based on the lowest Spearman’s rank correlation coefficient for the group (**A**), average Pearson correlation coefficients for the group (**B**) or the lowest Pearson correlation coefficients for the group (**C**). The resulting dendrograms (also see Figure 1C) are the same regardless of the approach. Datasets assigned to larger (80kb or 160kb) bins yield equivalent dendrograms, which are also robust to parameter changes. When chromatin contacts are binned to smaller segments (5kb, 10kb, 20kb), the clustering outcomes start to vary depending on the kind of correlation coefficient (Spearman or Pearson) and the grouping approach (lowest or average for the group) is used. We, therefore, consider the latter results unreliable.

**Figure S5.**
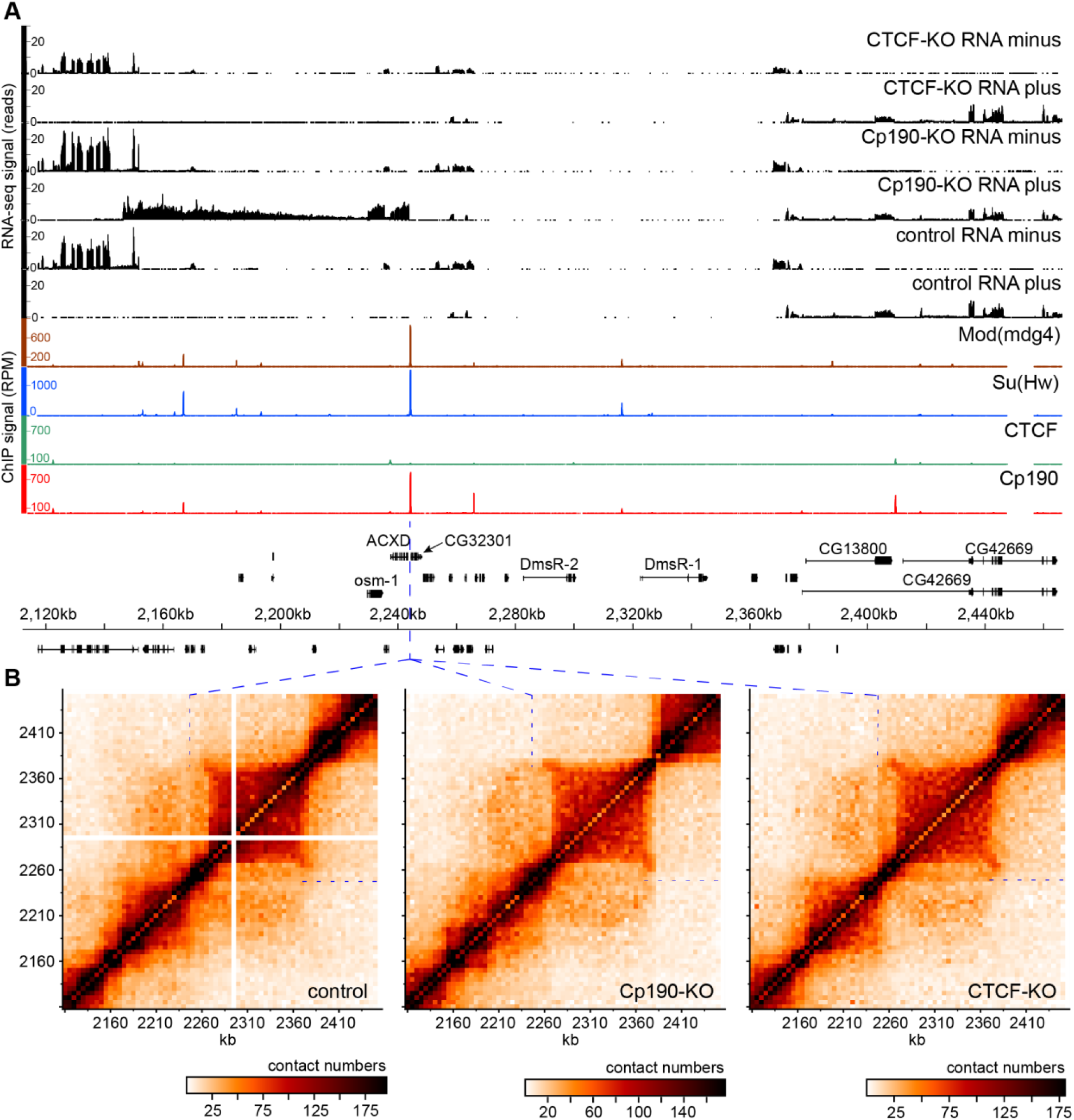
Chromatin topology around the *62D* insulator element. **A.** Genomic organization of the 62D region. ChIP-seq profiles for Cp190, CTCF, Su(Hw) and Mod(mdg4) proteins in control cells are displayed as number of sequencing reads per position per million of total reads. RNA-seq profiles from control, Cp190-KO and CTCF-KO cells are displayed separately for each DNA strand as number of sequencing reads per position. Positions and the exon structure of annotated transcripts are shown above (transcribed from left to right) or below (transcribed from right to left) the scale in *dm6*, 2014 genome release coordinates. The position of the *62D* insulator element is indicated with blue dashed lines. **B.** Chromatin contacts within the 62D region in control, Cp190-KO and CTCF-KO cells. Contacts measured by individual Hi-C experiments were assigned to 5kb bins and normalized by iterative correction (*89*). The data from replicate experiments were combined and visualized with the *gcMapExplorer* software (*80*).

**Figure S6.**
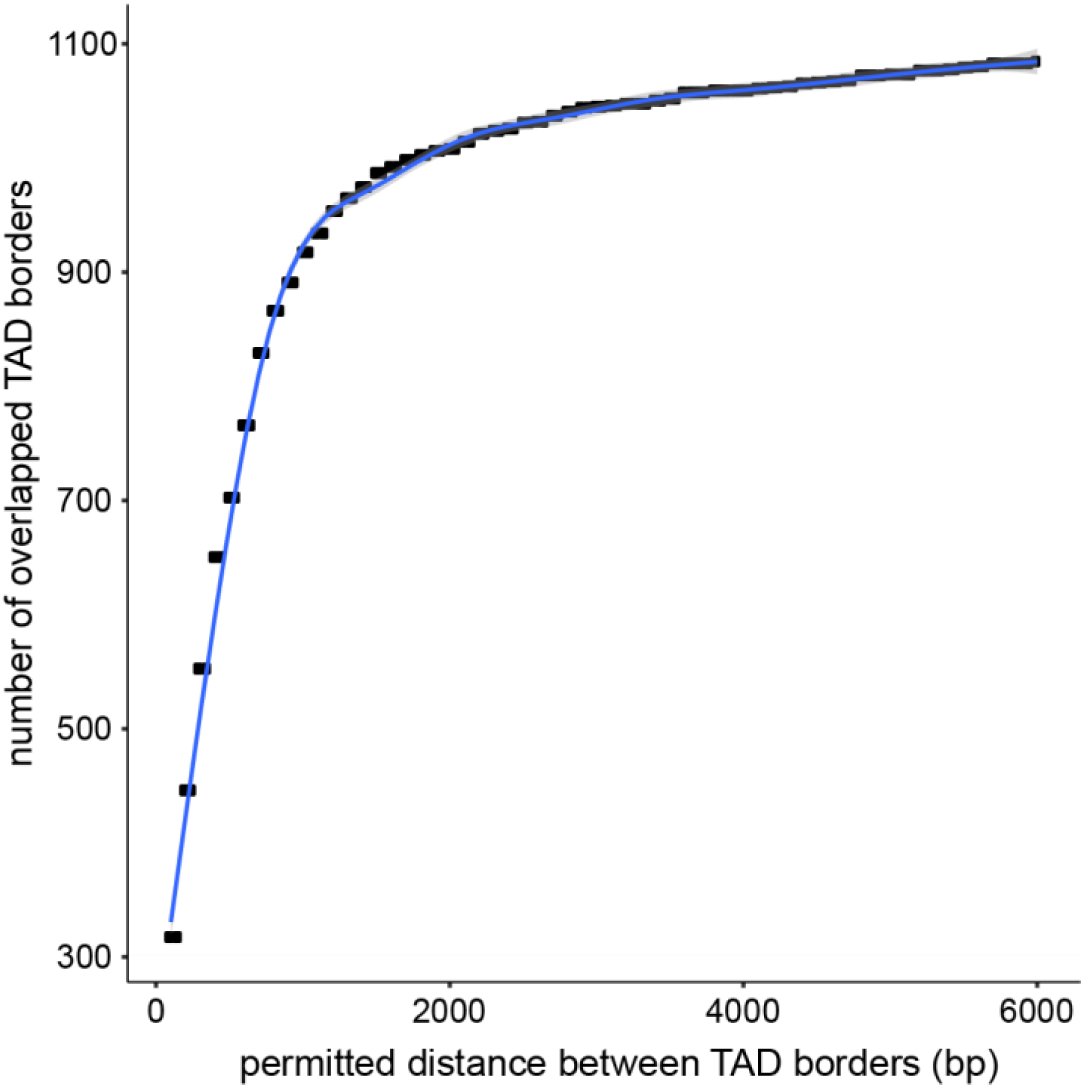
The optimal accuracy of TAD border mapping. To estimate the accuracy of TAD border positions the number of overlapped TAD borders defined in two replicate experiments in the control cells was plotted against the permitted distance between two borders considered overlapped. From this, TAD borders identified in replicate experiments at a distance of 2000bp or less were considered identical.

**Figure S7.**
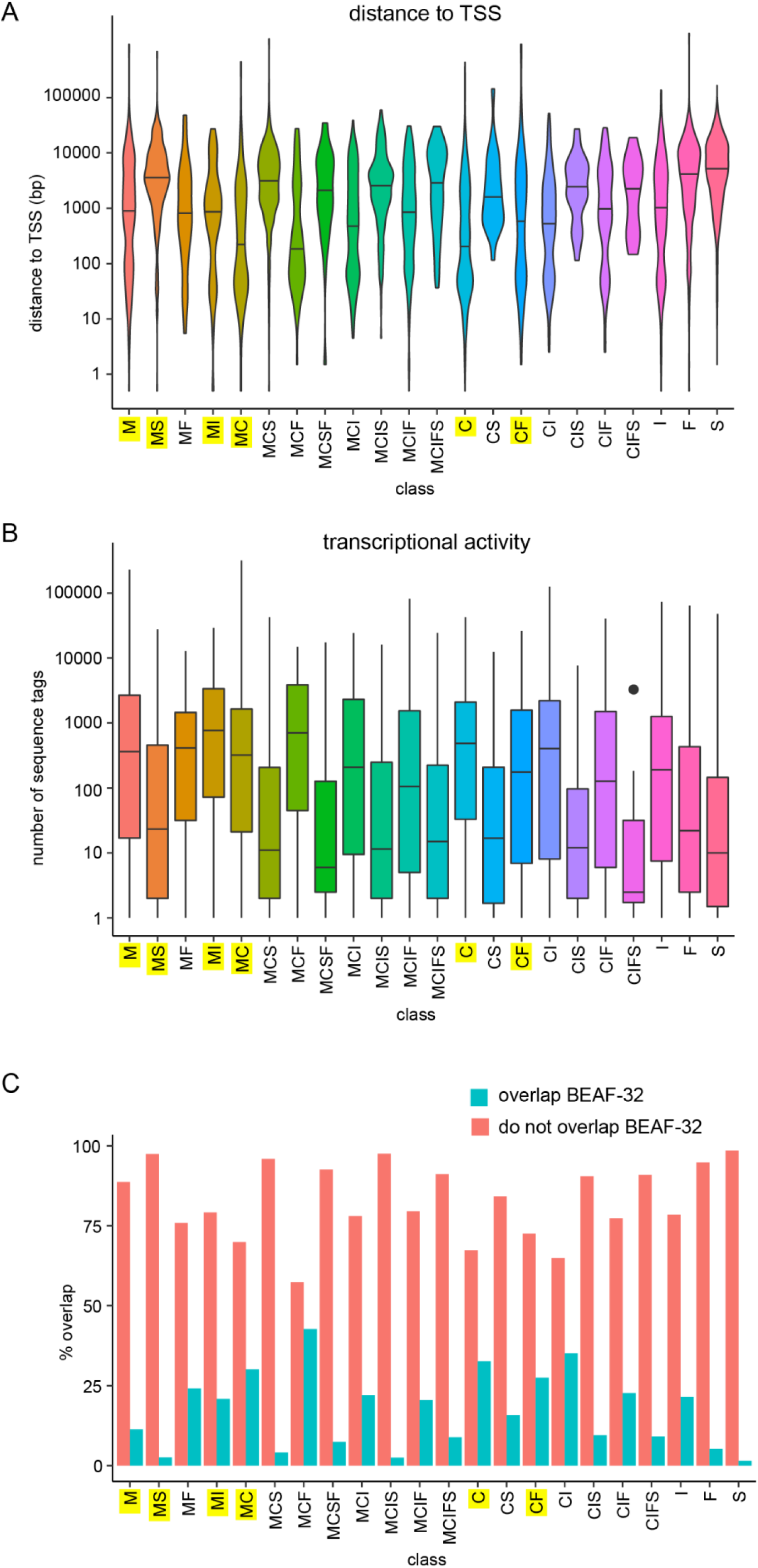
Relation between classes of insulator protein bound regions, proximity to transcription start sites, transcription and BEAF-32. **A.** Violin-plots show distances to the closest TSS for different classes of insulator protein bound regions. Horizontal lines indicate medians. Here and in **B** and **C**, the yellow shading marks the classes discussed in the text. **B.** Transcriptional activity of the gene with the closest TSS for different classes of insulator protein bound regions. Box-plots indicate the median and span interquartile range with whiskers extending 1.5 times the range and outliers shown as black dots. **C**. Bar-plots indicate fraction of insulator protein bound regions of different classes that overlap with BEAF-32 bound regions mapped by modENCODE (*37*).

**Figure S8.**
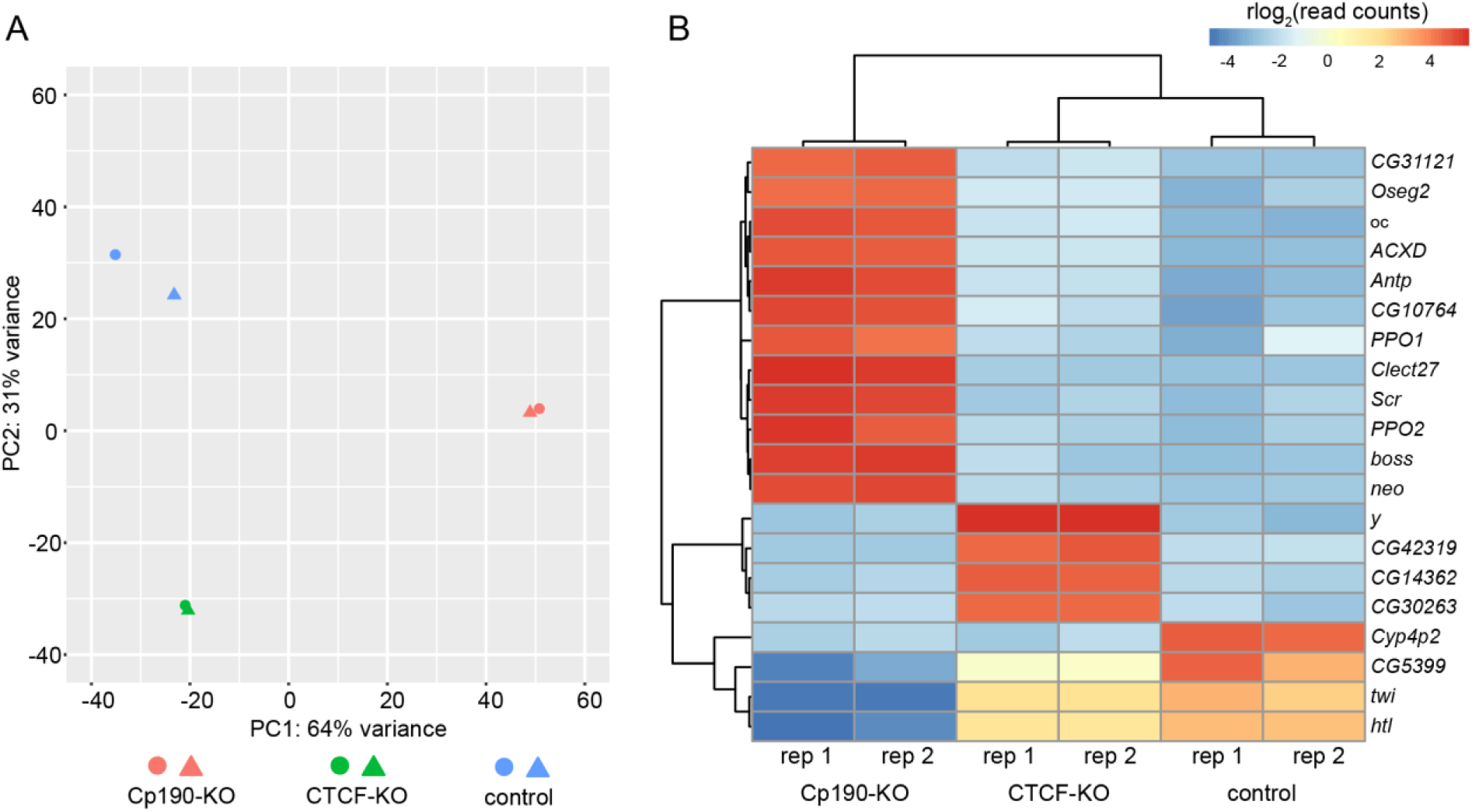
Differences in transcription between Cp190-KO, CTCF-KO and control cells. **A.** Principal component analysis of replicate RNA-seq experiments in the three cell lines. **B.** Clustered heatmap of RNA-seq signals for the twenty most differentially transcribed genes. Both analyses indicate that experiments are highly reproducible and that Cp190-KO is the most different of the three cell lines.

**Figure S9.**
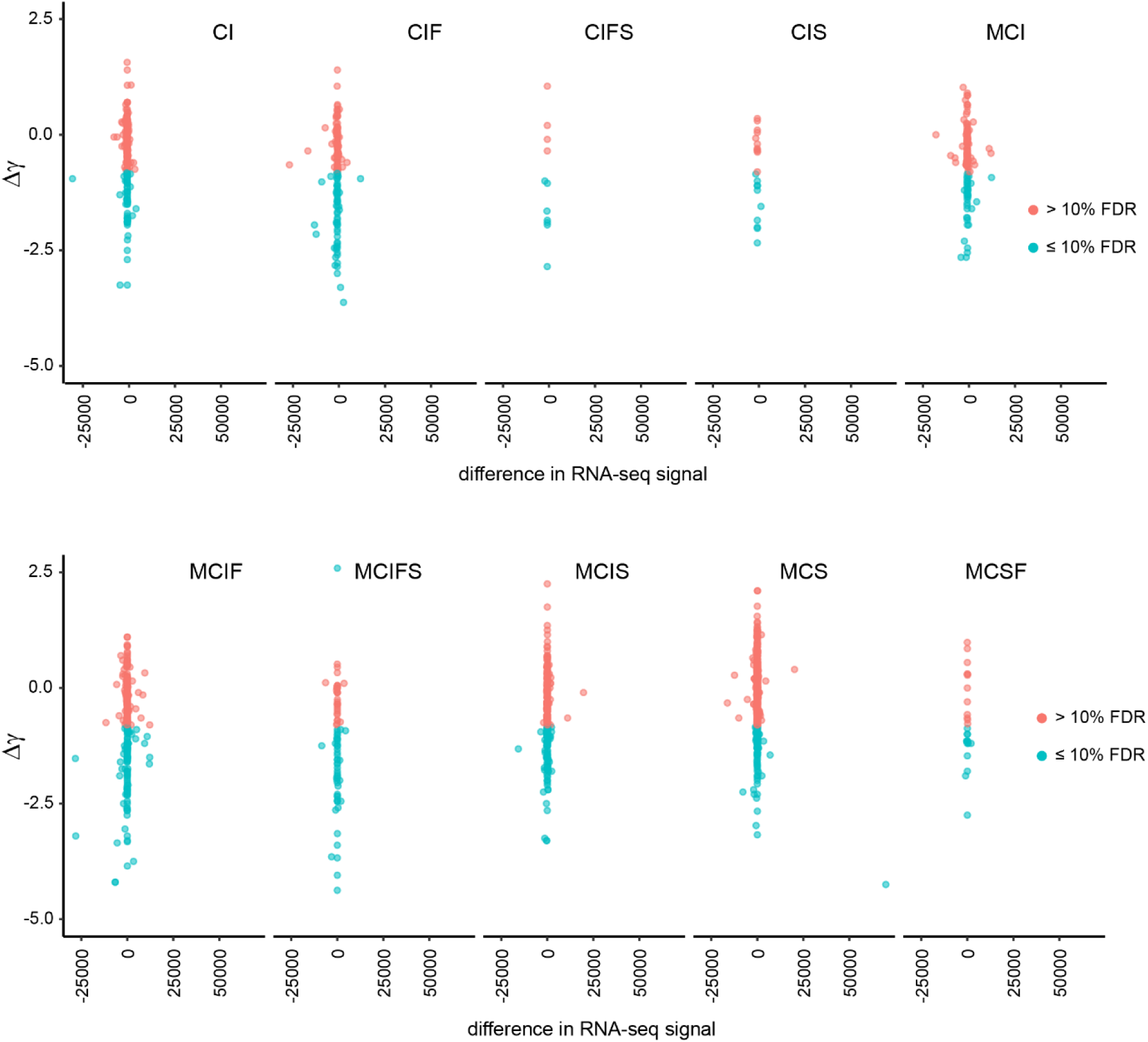
No correlation between changes in distance scaling factors (Δγ) and transcription of the nearby genes. Scatter-plots compare the changes in distance scaling factors (Δγ) at insulator protein binding sites of various classes to differences in RNA-seq signals of genes closest to these sites after Cp190 knock-out.

**Figure S10.**
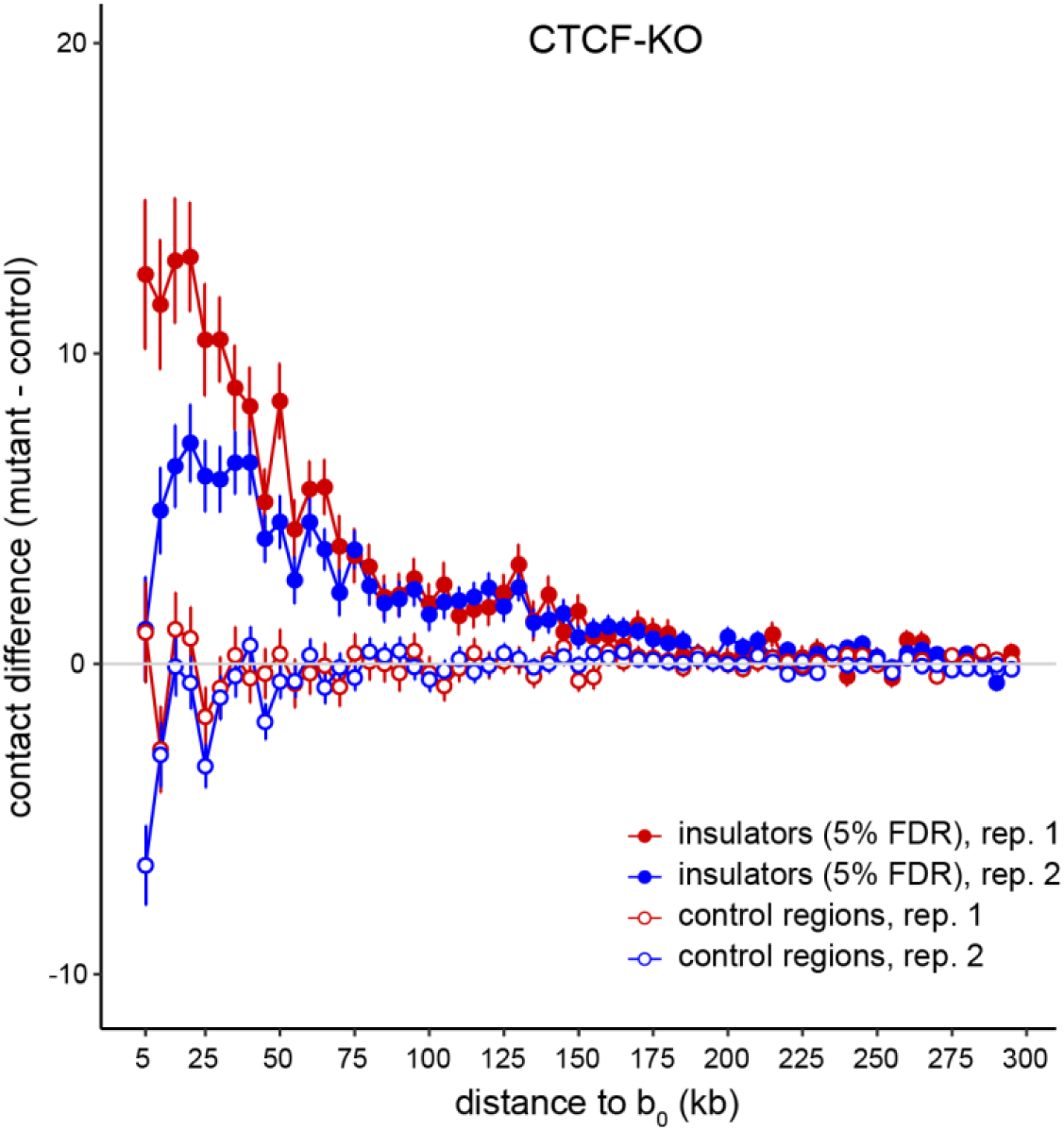
Action range of CTCF-dependent insulators. Average contact crossing difference curves for CTCF-dependent insulator elements (filled circles) and randomly selected control regions that do not bind any insulator proteins (empty circles) determined from two replicate Hi-C experiments (indicated with red and blue colours). Note that at close distances (5-10kb) the estimate of chromatin contact frequency from proximity ligation (the underlying principle of Hi-C method) becomes less reliable because, most of the time, chromatin fragments are sufficiently close to each other and the likelihood of successful ligation is more dependent on random chance.

