## Supplementary code for "Topological screen identifies hundreds of Cp190 and CTCF dependent *Drosophila* chromatin insulator elements": gen_ContactCcrossingCurves.sh

```
# generate pairwise contact

# IC map
bin_p="../../20190517_gcmap2bin-contact"
map_p="../../20210524_calculatePair/n_contactAftermultiHiCcompare"

# hicmaptools cmd
cmd="~/github_projs/hicmaptools/bin/hicmaptools"

bat_f="../sources/insulators_all_sorted.tsv"

# all HiC data
HiCs=( Ras3_rep1 Ras3_rep2 CTCF_rep1 CTCF_rep2 CPR6_rep1 CPR6_rep2 )
bin_dis=( 1 2 3 4 5 6 7 8 9 10 11 12 13 14 15 16 17 18 19 20 21 22 23 24 25 26 27 28 29 30 31 32 33 34 35 36 37 38 39 40 41 42 43 44 45 46 47 48 49 50 51 52 53 54 55 56 57 58 59 60 )

# loop HiC data
for HiC in "${HiCs[@]}"
do
	echo "${HiC}"
	for	dis in "${bin_dis[@]}"
	do
  		$cmd -in_map $map_p/${HiC}.binmap -in_bin $bin_p/${HiC}_5k.bins -bait $bat_f -ner_bin $dis -random 1 -output tsvs/${HiC}-n_${dis}.tsv &> logs/${HiC}-n_${dis}.log
  		rm tsvs/*.txt
  	done
done
```
