## Supplementary code for "Topological screen identifies hundreds of Cp190 and CTCF dependent *Drosophila* chromatin insulator elements": gen_InsulatorLoopingTest.sh

```
# generate pairwise contact

# IC map
bin_p="../../20190517_gcmap2bin-contact"
map_p="../n_contactAftermultiHiCcompare"

# hicmaptools cmd
cmd="~/github_projs/hicmaptools/bin/hicmaptools"

pair_f="../insulatorList/insulators2closePair.tsv"

# all HiC data
HiCs=( Ras3_rep1 Ras3_rep2 CTCF_rep1 CTCF_rep2 CPR6_rep1 CPR6_rep2 )

# loop HiC data
for HiC in "${HiCs[@]}"
do
  echo "${HiC}"
  $cmd -in_map $map_p/${HiC}.binmap -in_bin $bin_p/${HiC}_5k.bins -couple $pair_f -random 1 -output ${HiC}.tsv
done
```
