## Supplementary code for "Topological screen identifies hundreds of Cp190 and CTCF dependent *Drosophila* chromatin insulator elements": hic_normalize.R

```
library(multiHiCcompare)

args=commandArgs(trailingOnly =TRUE)
filepath<-args[1] #path of file
filepath <- "1_gcmap2rds/"
#load raw data
ras3rep1<-readRDS(paste0(filepath,"Ras3_rep1_original.rds"))
ras3rep2<-readRDS(paste0(filepath,"Ras3_rep2_original.rds"))
ctcfrep1<-readRDS(paste0(filepath,"CTCF_rep1_original.rds"))
ctcfrep2<-readRDS(paste0(filepath,"CTCF_rep2_original.rds"))
cpr6rep1<-readRDS(paste0(filepath,"CPR6_rep1_original.rds"))
cpr6rep2<-readRDS(paste0(filepath,"CPR6_rep2_original.rds"))

# make the hicexp object
groups <- factor(c(1, 1, 2, 2, 3, 3))
covariates <- data.frame(enzyme = factor(c('dpnii', 'dpnii','dpnii', 'dpnii','dpnii', 'dpnii')), 
						 batch = c(1, 2, 1, 2, 1, 2))
#hicexp <- make_hicexp(ras3rep1,ras3rep2,ctcfrep1,ctcfrep2,cpr6rep1,cpr6rep2,
#					  groups = groups,covariates = covariates,
#					  filter = F, remove_zeros = F, remove.regions=NULL)
hic_table(hicexp)

#save as rds file
saveRDS(hicexp,paste0(filepath,"rawhicdata.rds"))

#normalize
normal<-fastlo(hicexp)
hic_table(normal)

saveRDS(normal,paste0(filepath,"rawhicdata_normalize.rds"))
```
